# Ku70 suppresses alternative end-joining in G1-arrested progenitor B cells

**DOI:** 10.1101/2021.02.20.432121

**Authors:** Zhuoyi Liang, Vipul Kumar, Marie Le Bouteiller, Jeffrey Zurita, Josefin Kenrick, Sherry G. Lin, Jiangman Lou, Jianqiao Hu, Adam Yongxin Ye, Cristian Boboila, Frederick W. Alt, Richard L. Frock

**Affiliations:** Howard Hughes Medical Institute, Program in Cellular and Molecular Medicine, Boston Children’s Hospital, Department of Genetics, and Department of Pediatrics, Harvard Medical School, Boston, MA 02115, USA; Stanford University School of Medicine, Division of Radiation and Cancer Biology, Department of Radiation Oncology, Stanford, CA 94305, USA

**Keywords:** End-Joining, DSB Repair, Cas9, G1-phase, V(D)J Recombination

## Abstract

Classical nonhomologous end-joining (C-NHEJ) repairs DNA double-stranded breaks (DSBs) throughout interphase but predominates in G1-phase when homologous recombination is unavailable. Complexes containing the Ku70/80 (“Ku”) and XRCC4/Ligase IV (Lig4) core C-NHEJ factors are required, respectively, for sensing and joining DSBs. While XRCC4/Ligase IV are absolutely required for joining RAG1/2-endonucease (“RAG”)-initiated DSBs during V(D)J recombination in G1-phase progenitor lymphocytes, cycling cells deficient for XRCC4/Ligase IV also can join chromosomal DSBs by alternative end-joining (A-EJ) pathways. Restriction of V(D)J recombination by XRCC4/Ligase IV-mediated joining has been attributed to RAG shepherding V(D)J DSBs exclusively into the C-NHEJ pathway. Here, we report that A-EJ of DSB ends generated by RAG1/2, Cas9:gRNA and Zinc finger endonucleases in Lig4-deficient G1-arrested progenitor B cell lines is suppressed by Ku. Thus, while diverse DSBs remain largely as free broken ends in Lig4-deficient G1-arrested progenitor B cells, deletion of Ku70 increases DSB rejoining and translocation levels to those observed in Ku70-deficient counterparts. Correspondingly, while RAG-initiated V(D)J DSB joining is abrogated in Lig4-deficient G1-arrested progenitor B cell lines, joining of RAG-generated DSBs in Ku70-deficient and Ku70/Lig4 double-deficient lines occurs through a translocation-like A-EJ mechanism. Thus, in G1-arrested, Lig4-deficient progenitor B cells are functionally end-joining suppressed due to Ku-dependent blockage of A-EJ, potentially, in association with G1-phase down-regulation of Ligase1. Finally, we suggest that differential impacts of Ku-deficiency versus Lig4-deficiency on V(D)J recombination, neuronal apoptosis, and embryonic development results from Ku-mediated inhibition of A-EJ in the G1 cell cycle phase in Lig4-defcient developing lymphocyte and neuronal cells.

**Significance Statement:** Alternative end-joining (A-EJ) is implicated in oncogenic translocations and mediating DNA double-strand break (DSB) repair in cycling cells when classical nonhomologous endjoining (C-NHEJ) factors of the C-NHEJ Ligase complex are absent. However, V(D)J recombination-associated DSBs that occur in G1 cell cycle-phase progenitor lymphocytes are joined exclusively by the C-NHEJ pathway. Until now, however, the overall mechanisms that join general DSBs in G1-phase progenitor B cells had not been fully elucidated. Here, we report that Ku, a core C-NHEJ double-strand break recognition complex, directs repair of a variety of different targeted DSBs towards C-NHEJ and suppresses A-EJ in G1-phase cells. We suggest this Ku activity explains how Ku-deficiency can rescue the neuronal development and embryonic lethality phenotype of Ligase 4-deficient mice.

## Introduction

DNA double-strand breaks (DSBs) arise from sources both intrinsic and extrinsic to the cell, and improper DSB repair can lead to genomic instability and oncogenic translocations. To resolve DSBs, mammalian cells largely use two major classes of repair pathways: classical nonhomologous end-joining (C-NHEJ), which is active throughout interphase, and homology directed repair (HDR), which is only active in S/G2 cell cycle phases (Scully et al., 2019; Chang et al., 2017). In the absence of C-NHEJ, cycling cells have been found to also join DSB by an alternative end-joining pathway or pathways (Chang et al., 2017)

Programmed, cell-intrinsic DSBs are generated during V(D)J recombination in developing B and T lymphocytes. V(D)J recombination assembles V, D, and J gene segments into variable region exons within antigen receptor loci of lymphocyte progenitors during the G1 cell cycle phase (Teng and Schatz, 2015). The RAG1/2 (RAG) endonuclease is recruited to a recombination center in antigen receptor loci (Teng and Schatz, 2015; Lin et al., 2018), where it binds recombination signal sequence (RSS) located adjacent to V, D, and J gene segments in one of its two active sites (Kim et al., 2015; Ru et al., 2015). Then, the single RSS-bound RAG linearly scans long-range distances of adjacent chromatin in the locus, presented by cohesin-mediated loop extrusion, for compatible RSSs with which to mediate cleavage (Hu et al., 2015; Zhao et al, 2016; Jain et al., 2018; Lin et al., 2018; Zhang et al., 2019b, Ba et al., 2020; Hill et al., 2020; Dai et al., 2021). Once two RSSs are appropriately paired, RAG cleaves between the two sets of RSSs and their coding ends to form RSS and coding-end (CE) DSB ends that are held in a post-cleavage synaptic complex (Schatz and Swanson 2011). Joining of cleaved RSS ends to each other and coding ends each other, respectively, is subsequently carried out exclusively by C-NHEJ (Alt et al., 2013; Kumar and Alt 2016), potentially due to RAG shepherding the broken ends specifically into the C-NHEJ pathway (Lee et al., 2004; Corneo et al., 2007). V(D)J recombination end-specific joining is unlike most chromosomal translocations or deletions (involving, for example, Cas9:gRNA or other types of DSBs) in which a given DSB-end can join to either DSB end of the other DSB (Chiarle et al., 2011; Frock et al., 2015).

C-NHEJ contains the “core” factors Ku70/Ku80 (Ku), which form the DSB recognition complex, as well as XRCC4/Ligase IV (Lig4), which forms the DSB ligation complex. Core C-NHEJ factors are necessary for joining CEs and RSS ends during V(D)J recombination in G1-phase developing lymphocyte progenitors; accordingly, mice deficient for core C-NHEJ factors exhibit a severe combined immune deficiency (SCID) due to defective repair during V(D)J recombination (Alt et al., 2013; Kumar and Alt 2016). However, Ku70-deficient mice can have a “leaky” SCID when compared to a complete SCID in XRCC4/Lig4-deficient mice, consistent with low level of V(D)J recombination-like joining in the absence of Ku (Gu et al., 1997). Deficiency for core C-NHEJ factors also leads to substantial p53-dependent apoptosis of newly generated post-mitotic neurons (Frank et al., 2000; Gao et al., 1998; 2000); however, the impact of Ku deficiency on neuronal apoptosis is not nearly as severe as that of XRCC4 or Ligase4 (Lig4) deficiency (Gu et al., 2000).

As defined in the context of core C-NHEJ deficiency, cycling mammalian cells can also access an alternative end-joining (A-EJ) pathway (or pathways) to relatively robustly join DSB ends generated via translocations or during Immunoglobulin heavy chain (IgH) Class Switch Recombination (CSR) in mature B cells (Zhu et al., 2002; Yan et al., 2007; Boboila et al. 2010b). A-EJ also joins chromosomally I-SceI-generated DSBs in cycling mammalian cell lines (Guirouilh-Barbat et al., 2007; Boboila et al., 2012), fuses dysfunctional telomeres (Sfeir and de Lange et al., 2012), and promotes translocations to replication stress-enhanced recurrent DSB clusters in neural stem and progenitor cells (Wei et al., 2016). Implicated A-EJ factors include: Parp1, XRCC1/Ligase III (Audebert et al., 2004; Sfeir and de Lange, 2012), Pol θ (Mateos-Gomez et al., 2015; Ceccaldi et al., 2015), and RAD52 (Zan et al., 2017). Recent studies have also implicated Pol θ as specifically involved in A-EJ in the S/G2 phase (Yu et al., 2020). Cumulatively, studies of A-EJ have not fully addressed all contexts through which cells commit to A-EJ versus C-NHEJ (Mansour et al, 2013; Truong et al., 2013; Soni et al., 2014; Shamanna et al., 2016; Bakr et al., 2016; Kang and Yan 2018; Yu et al., 2020). In the context of V(D)J recombination, the post-synaptic RAG complex itself has been implicated in shepherding V(D)J RSS and coding end DSBs into the C-NHEJ versus A-EJ pathways in G1-phase progenitor B cell lines (Lee et al., 2004; Corneo et al., 2007; Kumar and Alt 2016). However, whether RAG generated DSBs can translocate to more general DSBs or whether repair of general DSBs in G1-arrested progenitor B cells can employ A-EJ have remained to be determined (Yu et al, 2020).

XRCC4 or Lig4-deficient mice succumb to embryonic lethality, which along with their severe neuronal apoptosis is rescued by p53-deficiency, with rescue of neuronal development, and embryonic development potentially occurring by rescue of newly generated Lig4 and XRCC4-deficient neurons from p53-dependent apoptosis in the presence of large numbers of unrepaired DSBs (Frank et al., 2000; Gao et al., 2000). In contrast, Ku-deficient mice do not have an embryonic lethal phenotype and, correspondingly, exhibit a much milder neuronal apoptosis phenotype (Nussenzweig et al.,1996; Gu et al., 1997; Gu et al., 2000). Notably, Ku-deficiency rescues the embryonic lethality of Lig4-deficiency mice and has related effects in cell lines (Adachi et al., 2001; Karanjawala et al., 2002). In this context, Ku binding to unrepaired breaks in the context of Lig4- or XRCC4-deficient newly generated neurons has been speculated to suppress their ability to repair persistent DSBs by A-EJ and, thereby, promote their survival (Haber, 2008; Alt et al. 2013). As DSBs can be substantially joined by A-EJ in cycling Lig4- or XRCC4-deficient cells (Yan et al., 2007; Guirouilh-Barbat et al., 2007), such Ku-dependent down-regulation of A-EJ could in theory have more impact in non-cycling cells such as neurons and G1-phase progenitor B cells.

To ascertain whether additional mechanisms that might restrict joining of RAG-initiated DSBs and determine whether joining of other types of DSBs is also restricted in the G1 cell cycle phase, we mapped RAG-, Cas9:gRNA- and Zinc finger nuclease-generated DSB repair fates in G1-arrested *v*-*Abl*-transformed progenitor B cell lines through a version of our linear amplification-mediated, high-throughput, genome-wide translocation sequencing (LAM-HTGTS) (Hu et al., 2016) modified to map bait DSB rejoining at single nucleotide resolution.

## Results

### Repair of Cas9:gRNA DSBs in G1

We derived two independent clonal wildtype v-*Abl*-transformed progenitor B cell lines (Abl lines) designated “WT A” and “WT B” Abl lines, respectively (see Methods). We derived clonal “Ku70^-/-^A1” and “Lig4^-/-^A1” Abl lines from WT A and “Ku70^-/-^B1” and “Lig4^-/-^B1” from WT B Abl lines, respectively, (see Methods). We also obtained two independent Lig4-deficient (Lig4^-/-^) Abl lines (Bredemeyer et al., 2006; Lee et al., 2013), which we termed “Lig4^-/-^C” and “Lig4^-/-^D” lines (Fig. S1A-G). As reported in earlier studies (Gu et al., 1997) loss of Ku70 decreases Ku80 protein to virtually undetectable levels, resulting in a complete loss of the Ku complex (Fig. S1A, S1E). Treatment with STI-571 for all Abl lines described causes G1-arrest, induction of RAG expression, and V(D)J recombination at endogenous antigen receptor loci including the Ig kappa (Igk) locus with Abl lines surviving under G1-arrest throughout experiments due to the Eμ-Bcl2 transgene present in the original A and B wildtype lines (Bredemeyer et al., 2006).

We introduced a Cas9:guide RNA (gRNA) bait DSB targeting the first intron of *c-Myc* (Hu et al., 2015) in G1-arrested Abl cells and employed HTGTS-JoinT-seq (HTGTS-based re<u>Join</u>ing and <u>T</u>ranslocation sequencing)(Supplement Methods) to measure genome-wide “prey” translocations from the 5’ “bait” DSB broken end and to measure imperfect rejoining of the 5’ bait DSB to its corresponding 3’ broken end, both at single nucleotide resolution. The use of a modified pipeline was necessary given that the original LAM-HTGTS translocation pipeline (Hu et al., 2016) detects only very few imperfectly rejoined bait DSB junctions in G1-arrested WT Abl cells due to limited resection in the G1 phase (Fig. S2A-C). We performed three independent replicates of experiments for each line assayed, for a combined total of approximately 100,000 junctions from each WT Abl line (Fig. 1A, B and Table S1). Genomewide plots from the Cas9:*c-Myc* bait for each G1-arrested, WT A and B Abl cells displayed three distinct junction groups: 1) bait DSB ends rejoined to each other (i.e. the bait break-site), 2) recurrent translocations to RAG-initiated DSBs in antigen receptor loci and 3) low-level, widespread translocations (Alt et al., 2013; Frock et al., 2015) (Fig. 1A, B). As predicted, the majority of recovered junctions from G1-arrested WT A and WT B Abl cells involved rejoining of the bait DSB ends (approximately 97% of total junctions) with a limited mean resection distance (<1bp) (Fig. 1A-E and Tables S1, S2). In this regard, rejoined junctions predominantly harbored small insertions (approximately 70-90%), consistent with known polymerases that work with C-NHEJ (Pryor et al., 2015; Gouge et al., 2015; Stinson et al., 2020), with the remaining junctions being either direct joins (5-20%) or employing short microhomologies (MHs)(1-10%) (Fig. 1F, G). Given the very high proportion of bait DSB rejoins relative to translocations, we have focused here on rejoining outcomes of the bait DSB ends except for internal RAG-generated DSBs Igκ locus joining which occurs more frequently.

**Figure 1.**
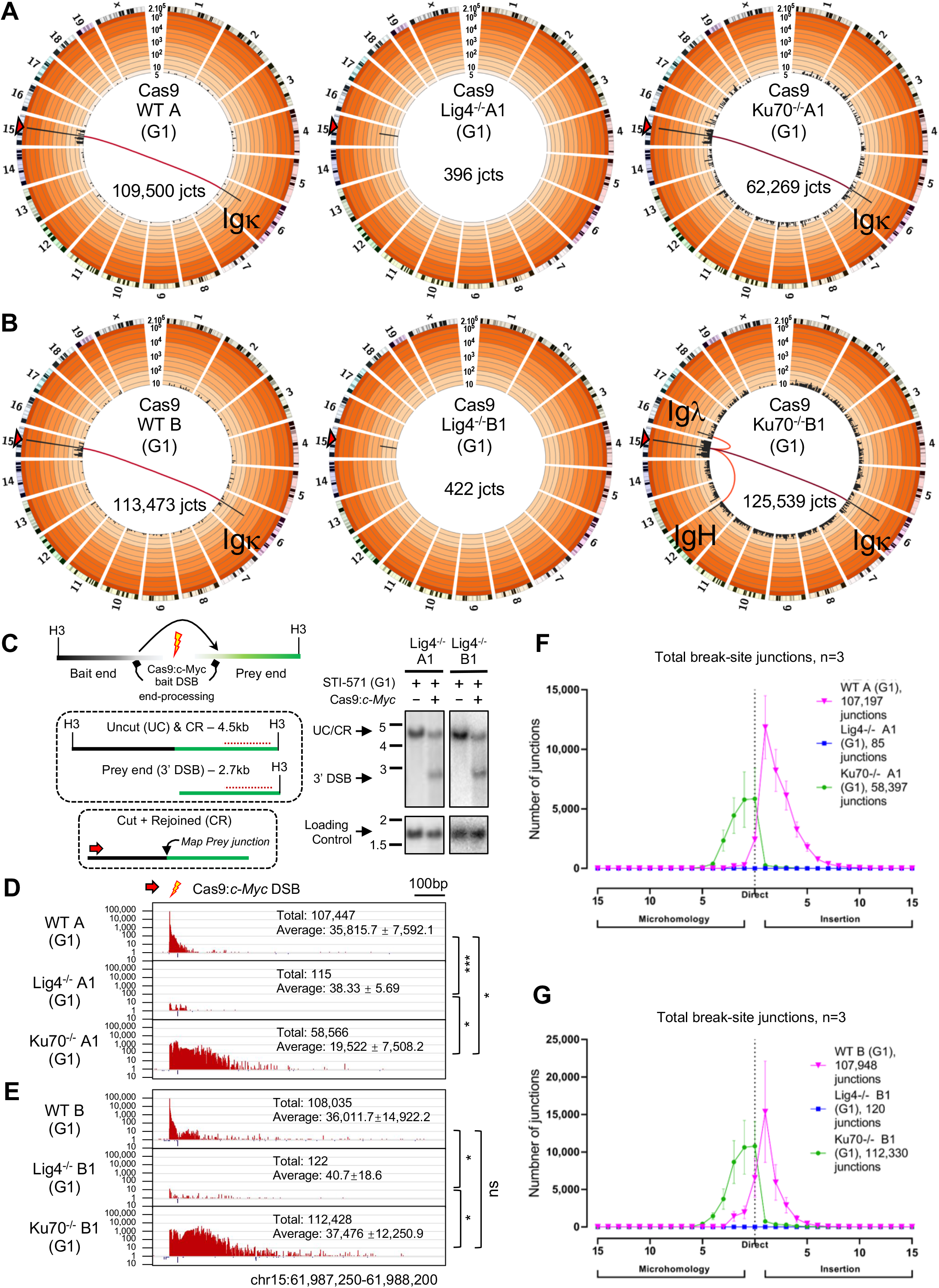
Disparate G1-arrested A-EJ outcomes of Cas9:*c-Myc* DSBs from specific core C-NHEJ deficiencies. A-B) Genome-wide prey-junctions from independently derived WT A and B, Lig4^-/-^A1 and B1 and Ku70^-/-^A1 and B1 Abl lines are binned into 5Mb regions (black bars) and plotted on Circos plots displaying a 1/2/5 increment log_10_ scale (Frock et al., 2015). Bar height indicates junction frequency. Frequency ranges are colored by order of magnitude from very light (<10) to dark orange (>10^5^). The light to dark red tone lines connecting the bait break to Ig loci represents translocation hotspots of greater significance. Circos plots are from pooled libraries after normalization, with total junctions indicated (*n*=3 for each clone; see Table S1). C) Left: diagram of Cas9:*c-Myc* bait DSB locus (top) with Southern fragments (H3 = HindIII) detected by probe (middle; red dashed line) and HTGTS-JoinT-seq emphasizing Cut + Rejoined junctions (lower; red arow = bait primer). Right: Southern for Lig4^-/-^ A1 and B1 clones; cutting and G1-arrest are indicated. D-E) Bait break-site profiles for clone set A (D) and B (E). Junctions in both (+)(red) and (-)(blue) chromosome orientations plotted on log_10_ scale from pooled libraries indicated by total and average junction numbers (*n*=3 for each clone, Mean±SD; One-way ANOVA with post-hoc Tukey’s test, *P*<0.05, *, *P*<0.001, ***, ns = not significant). F-G) Microhomology and insertion utilization of bait break-site junctions are plotted with indicated lengths (*n*=3 for each clone, Mean±SD).

Lig4^-/-^A1 and Lig4^-/-^B1 Abl cells were G1-arrested and then expressed with Cas9:*c-Myc* to measure repair in the absence of C-NHEJ. Strikingly, we recovered very few junctions with the total being 100-1,000 fold fewer than the number recovered from WT clones and approaching background levels (Fig. 1A-E and Table S1). Southern blotting analyses of the Cas9:*c-Myc* bait break-site revealed little change in the germline band, consistent with dominant rejoining with limited resection; however, a substantial fraction of the *c-Myc* alleles in G1-arrested Lig4^-/-^ Abl cells were present as un-joined DSB ends (Fig. 1C and Fig. S3A-C), demonstrating substantial bait site cutting but little rejoining in the latter. Retroviral complementation of Lig4^-/-^A1 Abl cells with Lig4 (Fig. S1B) restored joining patterns, junction structures, and median resection distance similar to those observed in WT (Fig. 2A-C and Tables S1, S2) and, correspondingly, with a decreased level of Cas9:*c-Myc* bait broken ends as detected by Southern blotting (Fig. S3D.) Finally, cycling WT A and Lig4^-/-^A1 Abl cells robustly rejoined the Cas9:*c-Myc* bait DSB ends. The majority of junctions from cycling WT A harbored short insertions over a mean resection distance of approximately 70bp, whereas cycling Lig4^-/-^ A1 were nearly all short MHs over a mean resection distance of approximately 300bp (Fig. S4A, B, Tables S1, S2), and no discernable change was observed in the Southern blot band harboring Cas9:*c-Myc* bait DSB (Figs. S4C). We conclude that the inability to rejoin in G1-arrested Lig4^-/-^ Abl cells is specific to both the absence of Lig4 and the G1 cell cycle phase.

**Figure 2.**
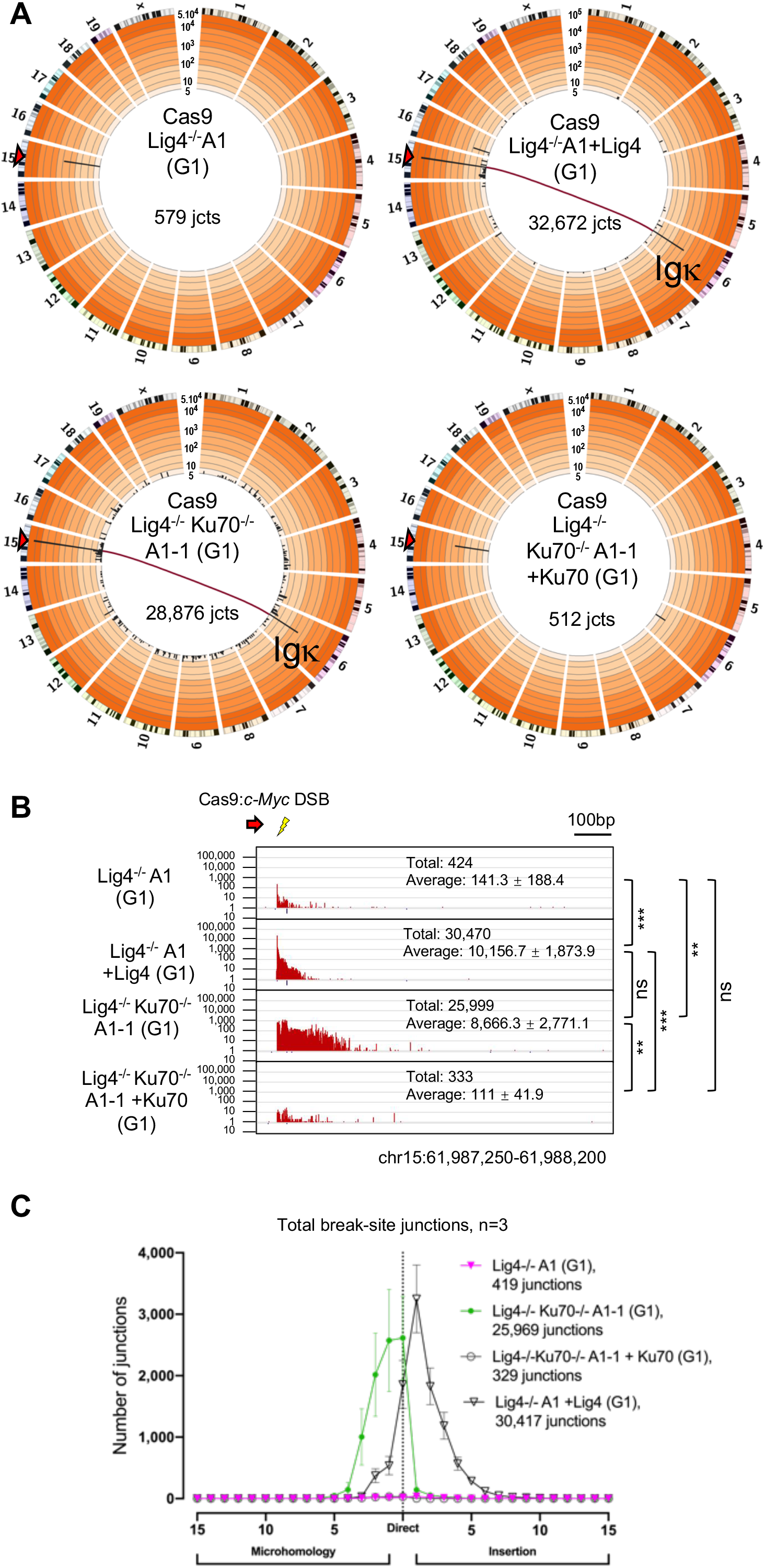
Ku70 suppresses the rejoining and translocation of Cas9:*c-Myc* DSBs in Lig4-deficient G1-arrested Abl cells. A) Genome-wide prey junctions from Cas9:*c-Myc* bait DSBs displayed as Circos plots (see Figure 1 legend; Table S1) for G1-arrested Lig4^-/-^A1, Lig4^-/-^A1 +Lig4, Lig4^-/-^Ku70^-/-^A1-1, and Lig4^-/-^Ku70^-/-^A1-1 +Ku70 Abl cells. B) Bait break-site junction profiles for Abl lines described in (A). Junctions are plotted similar to Fig. 1D (*n*=3 for each clone; One-way ANOVA with post-hoc Tukey’s test, *P*<0.01, **, *P*<0.001, ***; see Figure 1 legend). C) Microhomology and insertion usage of break-site junctions are plotted with indicated lengths (*n*=3 for each clone, Mean±SD) for the Abl lines and conditions described in (A).

Ku-deficiency has differential impacts on organismal development than XRCC4/Lig4-deficiency (Gu et al., 1997; Frank et al., 1998). Thus, we evaluated A-EJ of Cas9:*c-Myc* DSBs in G1-arrested Ku70^-/-^A1 and B1 Abl cells. Contrary to the near complete absence of A-EJ in G1-arrested Lig4^-/-^ Abl cells, Ku-deficient A-EJ was robust with recovered junctions at similar to slightly lower levels than that of WT Abl cells (Fig. 1A, B and Table S1, S2). Notably, Ku deficiency increased the mean resection distance of bait DSB rejoined junctions (to approximately 45bp) compared to WT Abl cells (Table S2). Despite the robust G1-arrested Ku70^-/-^ A-EJ (Fig. 1D, E), Southern blotting analysis displayed substantial unrepaired bait DSB ends (Fig. S3A-C). Ku-deficient rejoined junction structures predominantly consisted of small MHs (approximately 66%), but with a high fraction of direct joins (approximately 30%)(Fig. 1F, G).

Because Ku deficiency can rescue the embryonic lethality of Lig4 deficiency (Karanjawala et al., 2002), we hypothesized that the presence of Ku in G1-arrested Lig4^-/-^ Abl cells may block access of DSBs to A-EJ (Alt et al., 2013). To test this, we deleted Ku70 from Lig4^-/-^A1 and C Abl cell lines (generating Lig4^-/-^Ku70^-/-^A1-1 and C1 lines)(Fig. S1A, E) and assayed rejoining outcomes. Indeed, loss of Ku70 in Lig4^-/-^ Abl cell lines fully restored G1-phase joining patterns (e.g. increased resection, rejoining of bait ends, and direct junction utilization) to those observed in G1-arrested Ku70^-/-^A1 and B1 Abl cells (Fig. 2A-C and Figs. S5A, B, S6A-C, Tables S1, S2). Furthermore, complementation of Ku70 in Lig4^-/-^Ku70^-/-^ A1-1 and C1 Abl cell lines (Fig. S1E), suppressed repair levels approximately 100-fold, to low levels that approached those of G1-arrested Lig4^-/-^ Abl cell lines (Fig. 2A-C and Figs. S5A, B, S6A-C, Table S1), demonstrating that rejoining suppression was indeed Ku70-specific. We conclude that in G1-arrested Lig4^-/-^ Abl cells, A-EJ for Cas9 DSBs is suppressed by the Ku C-NHEJ DSB recognition complex.

### Repair of 5’ overhangs in G1

To determine whether DSB repair differences between C-NHEJ and A-EJ proficient backgrounds extend beyond repair of blunt DSB ends, we employed Lig4^-/-^D Abl cells, which harbor a doxycycline inducible zinc finger nuclease (ZFN) that targets the TCRβ locus enhancer on chromosome 6 (Eb-ZFN)(Lee et al., 2013). Upon doxycycline treatment, the ZFNs generate DSBs containing 4bp long 5’ overhang ends (Smith et al., 2000). We then generated a WT equivalent control by retroviral expression of Lig4 in these Lig4^-/-^D Abl cells (Figs. S1F). G1-arrested Lig4^-/-^D Abl cells displayed approximately 20-fold fewer total junctions from Eb-ZFN-induced DSBs than their Lig4-complemented G1-arrested counterparts (Fig. 3A, B), and, unlike their Lig4-complemented G1-arrested counterparts, accumulated substantial levels of unrepaired broken ends (Fig. S7A, B). Notably, although at low levels, significant numbers of junctions from Eb-ZFN expressing G1-arrested Lig4^-/-^D Abl cells were recovered (Fig 1A-E, 3A, B and Table S1). Furthermore, junctions recovered from the G1-arrested Lig4-complemented Lig4^-/-^D abl cells again predominantly contained short insertions with a limited mean resection distance (5bp) similar to rejoined Cas9:*c-Myc* bait DSB ends in WT Abl cells (Fig. 3C and Table S1, S2).

**Figure 3.**
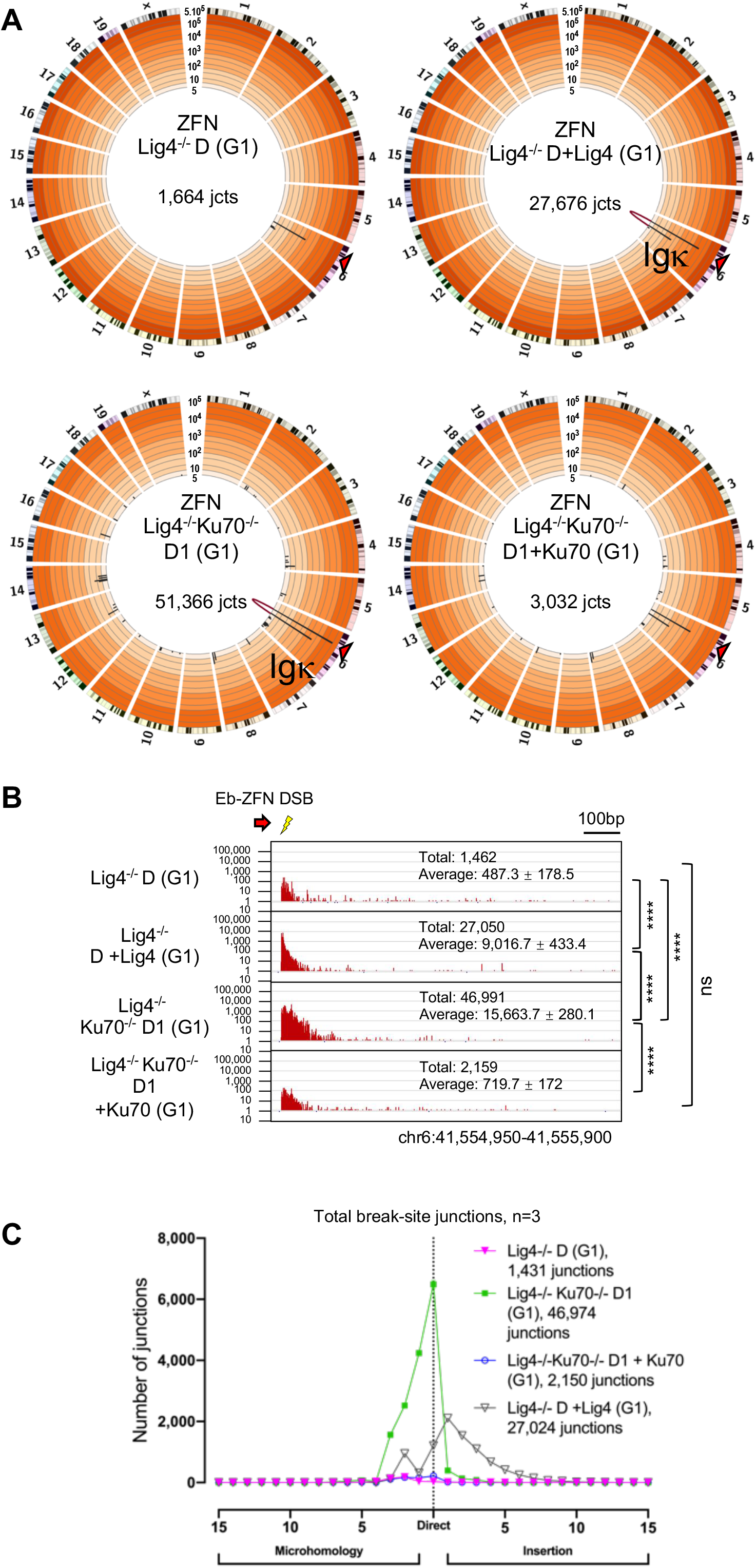
A-EJ of short 5’ overhangs from ZFN cleavage are suppressed by Ku in Lig4-deficient G1-arrested Abl cells. A) Genome-wide prey junctions from doxycycline-induced Eb-ZFN Lig4^-/-^D, Lig4^-/-^ +Lig4, Lig4^-/-^Ku70^-/-^ D1, and Lig4^-/-^Ku70^-/-^ D1 +Ku70 Abl cells displayed on Circos plots (see Figure 1 legend; Table S1). B) Induced Eb-ZFN bait break-site junction profiles from the G1-arrested Abl lines described in (A)(*n*=3 for each clone; One-way ANOVA with post-hoc Tukey’s test, *P*<0.0001, ****; see Figure 1 legend). C) Microhomology and insertion usage of Eb-ZFN break-site junctions from the Abl lines described in (A) are plotted with indicated lengths (*n*=3 for each clone, data represents Mean±SD).

To examine the influence of Ku on Eb-ZFN-generated DSBs, we deleted Ku70 from two separate clones of Lig4^-/-^D Abl cells to generate Lig4^-/-^Ku70^-/-^D1 and D2 Abl lines, which led to an approximately 20-30 fold increase in Eb-ZFN junctions recovered in the presence of Eb-ZFN; in this context, rejoined bait DSB junction structures predominantly contained short MHs (70%) but also had substantial numbers that were direct (30%), consistent with Cas9:*c-Myc* bait findings (Fig. 3A-C and Figs. S7C, D, S8A-C, Table S1). Correspondingly, ectopic expression of Ku70 in Lig4^-/-^Ku70^-/-^D1 and D2 Abl lines (Fig. S1G), in the presence of Eb-ZFN, decreased junction numbers to levels comparable to Lig4^-/-^D Abl cells (Fig. 3A, B and Figs. S7C, D, S8A-C, Table S1). Collectively, we conclude that the inability to repair DSBs in G1-arrested Lig4^-/-^ Abl cells, due to Ku suppression of A-EJ, also largely applies to ends with short 5’ overhangs.

### A-EJ of RAG-initiated DSBs

Given that Ku suppresses A-EJ for Cas9 and ZFN DSBs (above) and that Ku70^-/-^ mice can display a leaky SCID phenotype (Gu et al., 1997), we revisited the extent to which A-EJ participates in a V(D)J recombination-like process. Thus, we employed HTGTS-V(D)J-seq to measure the joining of the Jκ1 CE to other potential bait DSB ends which includes the approximately 130 Vκ gene segment CEs and their associated RSS ends (Hu et al., 2015; Zhao et al., 2016; Jain et al., 2018; Zhang et al., 2019b, Ba et al., 2020; Dai et al., 2021). Vκ gene segments in Igκ are either in deletional (blue bars) or inversional (red bars) recombination orientation relative to Jκ gene segments (Fig. 4A-D and SI Appendix Fig. S9A-D). As expected for the Jκ1 CE bait, WT A and B Abl libraries generated hundreds of thousands of junctions, displayed as highly orientation-biased joining to individual VκCEs, consistent with the V(D)J recombination mechanism (Fig. 4A, B–and Table S3)(Hu et al., 2015; Zhao et al., 2016; Jain et al., 2018; Lin et al., 2018; Zhang et al., 2019b, Chen et al., 2020; Ba et al., 2020; Dai et al., 2021). Most Vκ to Jκ V(D)J recombination-mediated joints were either direct or contained short insertions (Fig. 4E and Fig.S9E), consistent with high level TdT expression in these lines (Fig. S9G), which obviates use of dominant MHs between Vκ and Jκ junctional sequences (Komori et al, 1993). We found little to no RAG DSB junctions in Lig4^-/-^ A1, B1, and C Abl cell lines, and, correspondingly, these lines accumulated stable Jκ broken ends (Fig. 4A-D and Figs. S9A-D, S10A, B, S11A, B, Table S3). Lig4-complementation of Lig4^-/-^ A1 and C Abl lines attenuated levels of Jκ broken ends and restored normal Vκ to Jκ joining with orientation bias, and junction structures characteristic of normal V(D)J recombination (Fig. 4A, B, E and Fig. S9C-F, S10C, S11B, Table S3).

**Figure 4.**
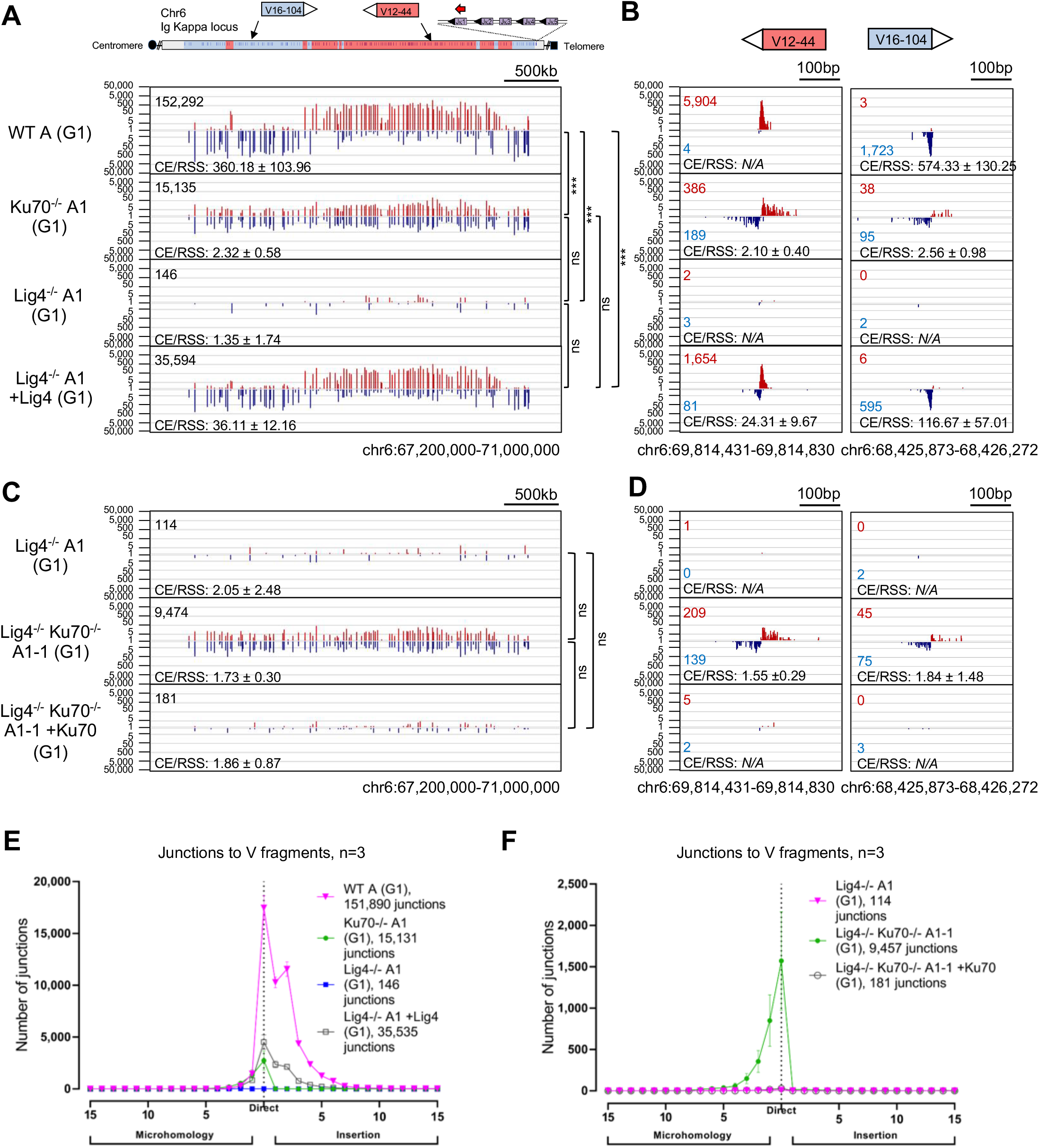
RAG DSBs are robustly joined to each other in the absence of the Ku DSB sensor by a translocation-based A-EJ mechanism. A) HTGTS-V(D)J-seq examining joining patterns of prey Vκ DSBs to the Jκ1 coding end comparing WT A, Ku70^-/-^A1, Lig4^-/-^A1 or Lig4^-/-^ A1 +Lig4 Abl cells. Blue bars indicate junctions positioned in the (-) chromosomal orientation and red bars indicate junctions in the (+) chromosomal orientation; depending on the relative orientation of the coding segments in Igκ, junctions of either orientation will represent joins to either CEs or RSSs generated from RAG cleavage at a Vκ RSS. Joins falling within 200bp of a CE or RSS are plotted (n=3 for each clone; One-way ANOVA to compare the ratios CE/RSS with post-hoc Tukey’s test, P<0.001, ***; see Table S4). B) Zoom-in of selected Vκ gene segments from (A). C) HTGTS-V(D)J-seq using the same approach as (A) but comparing Lig4^-/-^ A1, Lig4^-/-^Ku70^-/-^A1-1, and Lig4^-/-^Ku70^-/-^A1-1 +Ku70 Abl cells. D) Zoom-in of selected Vκ gene segments from (C). Microhomology and insertion usage in junctions joining from Jκ1 to Vκ segments are plotted with indicated lengths (*n*=3 for each clone, data represents Mean±SD).

In contrast to WT V(D)J recombination, we found significant levels of joining of the Jκ1 bait end to VκCEs and RSS ends in Ku70^-/-^A1 and B1 Abl cells, albeit at levels that were 5-10 fold lower than the levels of *bona fide* Vκ to Jκ junctions in WT A and B Abl lines (Fig. 4A, B and Fig. S9A, B, S10B, S11A, Table S3). Thus, the Jκ1 CE junctions in the Ku70^-/-^A1 and B1 Abl cells were not orientation-biased toward VκCEs like with C-NHEJ, but rather, were balanced for both VκCEs and VκRSSs (Fig 4A-D and Fig S9A-D, Table S3); thereby, indicating that they were joined by an A-EJ-mediated repair mechanism as opposed to end-specific joining by *bona fide* V(D)J recombination. A significant fraction of the Vκ to Jκ A-EJ mediated joins in Ku-deficient cells (approximately 30%) were potentially productive in that they fused the Vκ into the Jκ in frame (Fig. S9H) with a major subset of those that maintain Vκ and Jκ sequences with little or no deletion/resection similar to those of functional joins in WT cells (Fig. 4B). Finally, deletion of Ku in Lig4^-/-^A1 and C Abl cells rescued Vκ to Jκ junctions to levels comparable to those found in the context of Ku70-deficiency alone (Fig. 4C-F and Fig. S9C-F, S10D, S11C, Table S3). Such A-EJ mediated joins could be responsible for the leaky “V(D)J” recombination phenotype of Vκ to Jκ joining in Ku-deficient mice described previously (Gu et al., 1997).

## Discussion

We report that DNA DSBs generated by three different classes of endonucleases: RAG, Cas9:gRNA, and ZFN in G1-arrested Abl cells accumulate as un-joined broken ends in the absence of the Lig4 core C-NHEJ factor. We further define the mechanism as suppression of A-EJ in these G1-arrested cells by Ku. This finding is notable as it greatly contrasts with relatively robust A-EJ of general or CSR-associated DSBs reported previously in the absence of Lig4 or its essential XRCC4 co-factor complex in various cycling contexts (Boboila et al., 2010a; Boboila et al., 2010b; Boboila et al., 2012, Yu et al.,2020). While A-EJ is often considered a single pathway, two pathways of A-EJ have been proposed to be capable of mediating CSR in cycling B cells. One pathway, which occurs in Lig4-deficient B cells, uses almost exclusively MH-mediated joins and potentially represents a variation of C-NHEJ that likely uses Lig1 (Boboila et a., 2010b; Boboila et al., 2012). The second pathway found in the absence of either Ku or Ku plus Lig4, substantially uses direct joins, and represents a true A-EJ pathway, since it occurs in the absence of C-NHEJ recognition and joining components (Boboila et a., 2010b). The A-EJ pathway that operates in Ku-deficient G1-arrested Abl cells, generally matches well with the latter end-joining pathway and, thus, represents a *bona fide* A-EJ pathway. On the other hand, the C-NHEJ variant pathway that operates in CSR is essentially absent in G1-arrested Abl cells, consistent with very low level Lig1 expression in these and other G1-arrested cells (Fig. S12A-F, S13)(Akbari et al., 2009). Notably, however, joining of ZFN-generated DSBs in Lig4-deficient, G1-arrested Abl cells was substantially, but not completely suppressed by Ku, with recovered joins mostly MH-dependent, consistent with mediation by the variant C-NHEJ pathway using Lig3. There are various mechanisms by which ZFN-generated DSBs access this joining pathway at low-level, for example if ZFN-binding partially interferes with Ku-binding, but resolution of the mechanism will require further investigation.

Our findings provide a molecular mechanism that could enforce the shepherding of RAG-initiated DSBs to repair by C-NHEJ, versus A-EJ, during V(D)J recombination (Corneo et al., 2007; Deriano et al., 2011; Gigi et al., 2014; Lescale et al., 2016). Our findings also provide a cellular mechanism as to why Ku deficiency versus XRCC4/Lig4 deficiency differentially impacts developmental processes *in vivo*. With respect to lymphoid development, we have now mechanistically defined why low-level joining of RAG-initiated DSB joining occurs during V(D)J recombination in Ku70-deficient versus Lig4- or XRCC4-deficient mice (Gu et al., 1997; Gu et al., 2000). In this context, the inefficient translocation-based joining of Vκ and Jκ gene segments that escape the RAG post-synaptic complex are fused to make V(D)J-like joints of which a significant proportion can serve as functional joins (Hu et al., 2015). In addition, we speculate that our findings may explain the differential impact of Ku-deficiency versus Lig4- or XRCC4-deficiency on embryonic development. Thus, our findings support the hypothesis (Haber, 2008; Alt et al., 2013) that absence of Ku allows A-EJ to rejoin a substantial fraction of DSBs in postmitotic neurons and, thereby, does not impact embryonic viability; whereas, the abrogation of end-joining, mediated by the Ku complex, would lead to an intolerable DSB burden and embryonic lethality in the context of Lig4- or XRCC4-deficiency. This model also can explain why Ku-deficiency rescues embryonic lethality of XRCC4/Lig4-deficient mice (Karanjawala et al., 2002).

## Methods

### Generation of Cell lines and Reagents

“WT A” Abl cell line was generated from a Lig4^flox/flox^ Eμ-Bcl2 transgenic mouse as previously described (Boboila et al., 2010a). The “WT B” Abl cell line was generated from an Eμ-Bcl2 transgenic mouse. Lig4^-/-^ A1 was generated by Cre deletion, using adeno-Cre recombinase, from WT A. Ku70^-/-^ A1 was generated from WT A and Lig4^-/-^ Ku70^-/-^ A1-1 was generated from Lig4^-/-^ A1 using Cas9 (Table S4) Lig4^-/-^ B1 and Ku70^-/-^ B1 Abl lines were generated from WT B. Lig4^-/-^ C and Lig4^-/-^ D (Eb-ZFN) (Lee et al., 2013) were gifts from Barry Sleckman (University of Alabama at Birmingham); Lig4^-/-^Ku70^-/-^ C1 was derived from Lig4^-/-^ C; Lig4^-/-^Ku70^-/-^ D1 was derived from Lig4^-/-^ D (Table S4). All CRISPR/Cas9 generation and screening of knock-out cell lines was performed as previously described (Kumar et al., 2016). Abl cells were G1-arrested using STI-571 (3μM) for 96 hours (Supplement Methods).

### NGS sample preparation and HTGTS-JoinT-seq

Designer nuclease bait experiments employed HTGTS-JoinT-seq. Junction-enriched libraries were generated as previously described for LAM-HTGTS (Hu et al., 2016) but with the removal of the bait DSB rejoining blocking step. Input genomic DNA for each junction library sample/replication was 12μg and all sequences were aligned to the mm10 genome build. Libraries were sequenced by Illumina Miseq (250PE) or Nextseq (150PE) and primers specific to each are indicated (Table S4). Pooled raw sequences were demultiplexed and adapter trimmed using TranslocPreprocess (Hu et al., 2016). Libraries were then normalized to the same number of raw reads within each experiment. Translocations were identified using TranslocWrapper (Hu et a., 2016) with additional modifications, followed by mapping rejoined bait ends using the R module, JoinT (Supplement Methods). See SI Appendix for description of HTGTS-Rep-Rejoin.

### HTGTS-V(D)J-seq and Hotspot analysis

HTGTS-V(D)J-seq (Ba et al., 2020) was performed for the Jκ1 coding end bait as similarly described (Hu et al., 2015; Zhao et al., 2016) using the mm10 genome build and only junctions within 200bp of *bona fide* Vκ RSSs, with or without 150bp of *bona fide* Jκ RSSs were described. For all NGS experiments, MACS2-based algorithm was used with a false detection rate (FDR) cutoff of 10^-10 as described (Frock et al., 2015). Hotspots were called if enriched sites were significant in all replicate libraries.

### Southern analysis

Southern blots were performed with 20μg of input DNA of the given condition and digested with either HindIII (to detect unrepaired Cas9 DSB ends at *c-Myc* or ZFN DSB ends at Eb) or double digest with EcoRI and NcoI (to detect unrepaired Jκ DSB ends). Southern probes used and additional methods are indicated (Supplement Methods and Table S4).

### Data Deposition

Raw and processed sequencing data are available through GEO (GSE162453)

## Author Contributions

ZL, VK, RLF, FWA designed the study. ZL, VK, RLF, SGL performed experiments. JZ, VK, MLB, RLF developed HTGTS-Rep-Rejoin. MLB, RLF developed HTGTS-JoinT-seq. ZL, VK, MLB, JZ, JK, JH, AYY, FWA and RLF analyzed HTGTS data. JL, ZL analyzed GRO-seq data. CB generated a critical cell line. FWA, supervised research conducted at BCH/HMS, RLF supervised research conducted at Stanford University. ZL, VK, MLB, RLF, and FWA wrote the manuscript. Other authors helped polish the manuscript.

## Acknowledgements

We thank members of the F.W.A and R.L.F. laboratories for stimulating discussions and feedback. We thank Barry Sleckman (University of Alabama) for the Lig4^-/-^C and Eb-ZFN expressing Lig4^-/-^D Abl lines. This work was supported by NIH grant AI020047 to F.W.A., an NIH Ruth L. Kirchstein National Research Service Award Fellowships CA189740 to V.K. and AI117920 to S.G.L., Cancer Research Institute Training Grant to C.B., and a Radiation Research Foundation Career Development Award to R.L.F. F.W.A is an Investigator of the Howard Hughes Medical Institute. R.L.F. is a V Scholar for the V Foundation for Cancer Research.

