## Supplemental for "Ku70 suppresses alternative end-joining in G1-arrested progenitor B cells"

**Supplementary Information Methods**

**TranslocWrapper analysis pipeline update.** For all library analysis described, we used an updated component of the TranslocWrapper pipeline, TranslocDedup.R, which identifies duplicated junctions; the updated script now takes into account the nucleotide content of inserted sequences between the bait/prey junction. This updated version is available with HTGTS-JoinT-seq (link below) and can simply replace the same named script associated with TranslocWrapper (Hu et al., 2016).

**HTGTS-JoinT-seq.** Initial G1-arrest LAM-HTGTS experiments, modified to exclude the blocking step that would normally suppress rejoining of bait DSB ends, revealed a significant underrepresentation of bait rejoining using the TranslocWrapper pipeline (Hu et al., 2016)(Fig. S2A) which required appending a new bioinformatic rejoining module to work with TranslocWrapper in order to capture imperfect rejoining events along with translocations at single nucleotide resolution. We developed two independent bioinformatic DSB rejoining approaches, HTGTS-Rep-Rejoin (requiring HTGTS-Rep-seq)(Lin et al., 2016) and JoinT (using the same Bowtie2 aligner as with TranslocWrapper). While both approaches were in agreement with regard to similar magnitudes of junctions recovered for the HTGTS experiments performed in this study, JoinT accounted for more of the available sequence reads (Fig. S2A-C). Therefore, we described this adapted approach HTGTS-JoinT-seq (HTGTS-based reJoining and Translocation sequencing). When combined with TranslocWrapper, the rejoining module, JoinT, yielded ~3-1,000 fold more junctions, depending on the DSB repair background assayed.

**The JoinT module.** JoinT was developed in R and uses the standard result output of the TranslocWrapper translocation mapping pipeline (Hu et al., 2016), a common metadata file, and the fastq files of the libraries as arguments. JoinT specifically identifies single nucleotide junction alterations originating from the targeted bait DSB sequence and employs the open-source aligner, Bowtie2 (Langmead and Salzberg 2012) version 2.3.2, twice to find reads which align both to the bait (primer to DSB) and the downstream prey sequences (1.1kb or 10kb depending on G1 or cycling conditions; customizable) to collect alignment positions. Parameters for Bowtie2 are the same as with TranslocWrapper (Hu et al., 2016), with the exception of the minimum score (40) and seed length (-L) set to 8. Up to the 3 best scoring prey alignments were collected; the alignment with the highest proximity to the bait was chosen for further analysis. The resulting reads were then filtered to separate perfect bait/prey interface reads and modified junctions, based on the position of the end of the bait, the position of the start of the prey and the presence of insertions, if any. To enhance accuracy, the base call quality between the bait and the prey helped to distinguish between *bona fide* modifications of the sequence versus sequencing errors. Bait end rejoins were then combined with the translocations identified by TranslocWrapper. For each read found both as a translocation by TranslocWrapper and as a rejoin, JoinT kept the alignment with the highest proximity to the bait. To accommodate prey regions with repeated sequences, we evaluated alignments restricted to the repeated region. From these, we removed the alignments for which another alignment to the prey with a higher score was found and gave preference to the TranslocWrapper called junction if the alignment was superior; we used this option for the *c-Myc* locus (repeat in positions chr15: 61,987,339-61,987,386). Scripts and installation instructions for HTGTS-JoinT-seq can be located on Github: https://github.com/marielebouteiller/JoinT-seq

**The HTGTS-Rep-Rejoin module.** Rejoin, was adapted from HTGTS-Rep-seq (Lin et al., 2016) using the IgBlast aligner (Ye et al., 2013) and combines the identified rejoined data with translocations from the TranslocWrapper pipeline (Hu et al., 2016), requiring a common metadata file and the fastq files of the libraries. The python script, HTGTS-Rep, processed and stitched together raw paired end sequence reads and identified junctions using the bait sequence and the downstream prey sequence corresponding to the other side of the bait DSB as separate “gene segments” that IgBlast uses to discern joining configuration. The python script, Rejoin, then pulls junction information from HTGTS-Rep outputs to further filter for *bona fide* modifications versus sequencing/preparation artifacts by employing a 1nt step, 10bp sliding window along both bait/prey sequences to identify junction positions. Resulting junctions were then combined together with translocations identified separately using TranslocWrapper (Hu et al., 2016). Scripts, links to earlier HTGTS pipelines, and operating instructions for each bait rejoining detection approach are available here: https://github.com/marielebouteiller/JoinT-seq

**NGS analysis**. Genome-wide translocations are displayed using Circos plots (Frock et al., 2015). Regional profiles were generated using IGV (Robinson et al., 2011). GRO-seq was performed as described (Ba et al., 2020).

**Western blots**. Whole cell extracts from 48 hour cultured G1-arrested cells were generated under lysis conditions used previously (Kumar et al., 2016). Western blot for Lig4 and Ku80 was performed as previously described (Kumar et al., 2016). For Ku70 western blot, goat polyclonal sc-1486 (santa cruz, 1:500, overnight 4$^{o}$C probing in 5% milk) was used.

**Transfection**. At the time of assay, Abl pro-B lines were either asynchronously growing (Day 0), STI-571 G1-arrested (3µM) for 96 hours (Bredemeyer et al., 2006), or STI-571 G1-arrested for 48-60 hours and nucleofected with the Amaxa 4-D nucleofector (solution SF; program DN-100) according to manufacturer's instructions with the following modifications. For each clone, 45-60x10$^{6}$ cells were nucleofected at a ratio of 2µg pX330 Cas9 plasmid (targeting *c-Myc* (Hu et al., 2015))+100ng pMAX GFP (Lonza supplied)/3x10$^{6}$ cells. Nucleofected cells were cultured in STI-571-containing medium for the remainder of the experiment. Cells were harvested 48 hours after nucleofection and lysed for DNA as previously described (Kumar et al., 2016). Cells were harvested and collected for DNA in 2mL total lysis volume as previously described (Hu et al., 2016). Cycling Abl cells were nucleofected and harvested for genomic DNA as above. For ZFN cutting Abl pro-B lines, cells were arrested by 3µM STI-571 for 48 hours, then treated with 2µg/ml Doxycycline (Day 2, Sigma D9891) for another 48 hours before collecting for DNA as described above.

**Plasmids and retroviral infection**. Murine Ku70 coding sequence was amplified from a pool of cDNAs and cloned into pMSCV-puro-VK (Kumar et al., 2016). Lig4-pMSCV-puro was a gift from Kefei Yu (Michigan State University). Retroviral infection of Abl cells was performed as described (Kumar et al., 2016). The Cas9 pX330 plasmid (Addgene 42230) containing guide RNAs targeting *c-Myc* (Hu et al., 2015) or Ku70 loci (see Table S4) were generated as previously described (Cong et al., 2013).

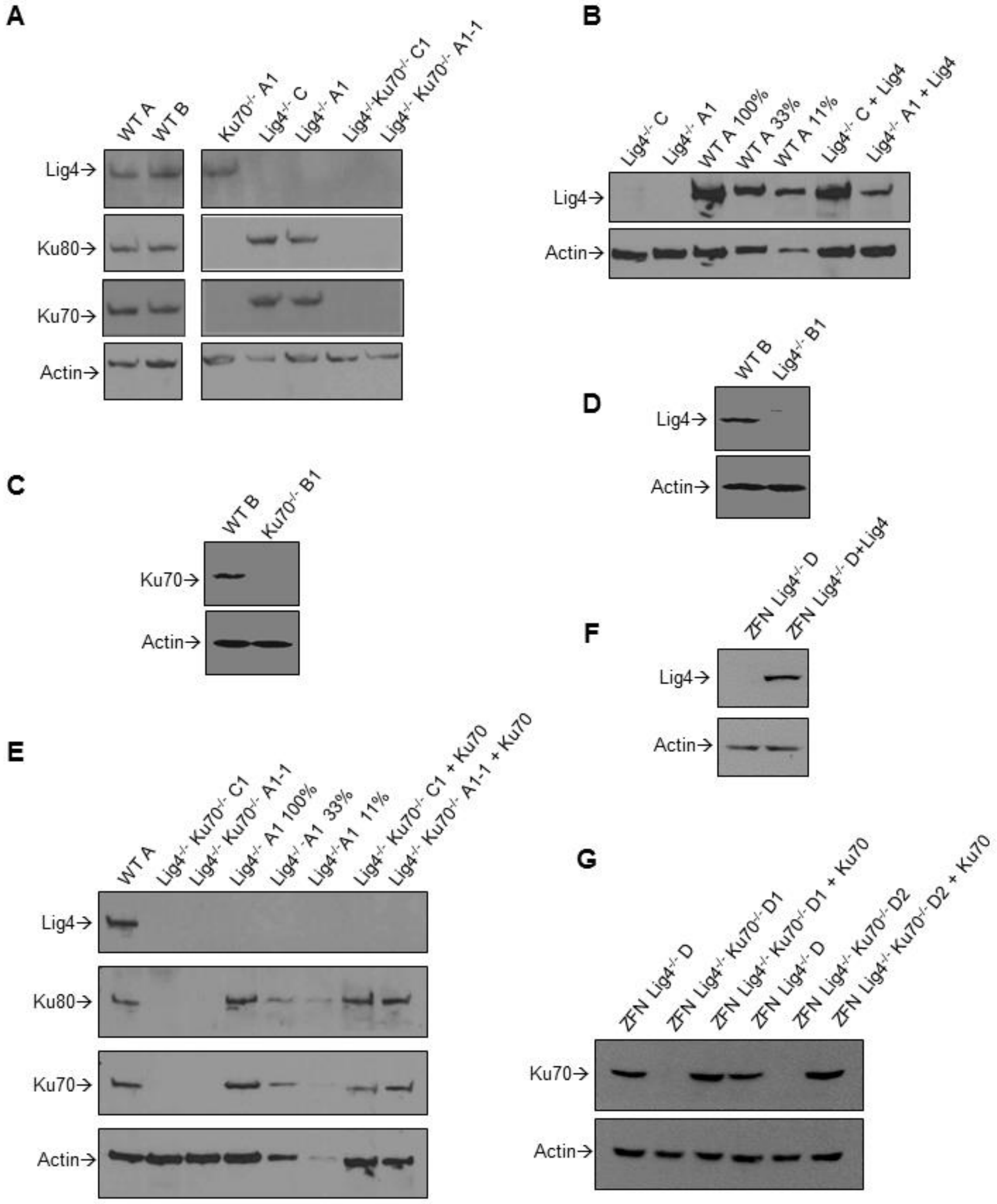


**Fig. S1. Western blot analysis of analyzed Abl cell lines.** A) Total protein levels of Lig4, Ku70, Ku80 in WT, Ku70$^{-/-}$, Lig4$^{-/-}$, or Lig4$^{-/-}$Ku70$^{-/-}$Abl cells measured by Western blot with indicated antibodies. Actin was the loading control. B) Lig4 complemented protein levels detected by Western blot for Lig4$^{-/-}$ Abl cells and compared to titrated WT Abl cells. C-D) Ku70 (C) and Lig4 (D) Western for Ku70$^{-/-}$B1 and Lig4$^{-/-}$B1 Abl clones. E) Ku70 complementation in Lig4$^{-/-}$Ku70$^{-/-}$A1-1 and C1 Abl cells with WT and titrated Lig4$^{-/-}$ controls. F-G) Lig4 (F) and Ku70 (G) complementation from Eb-ZFN Lig4$^{-/-}$D and Lig4$^{-/-}$Ku70$^{-/-}$D1 and D2 Abl cells.

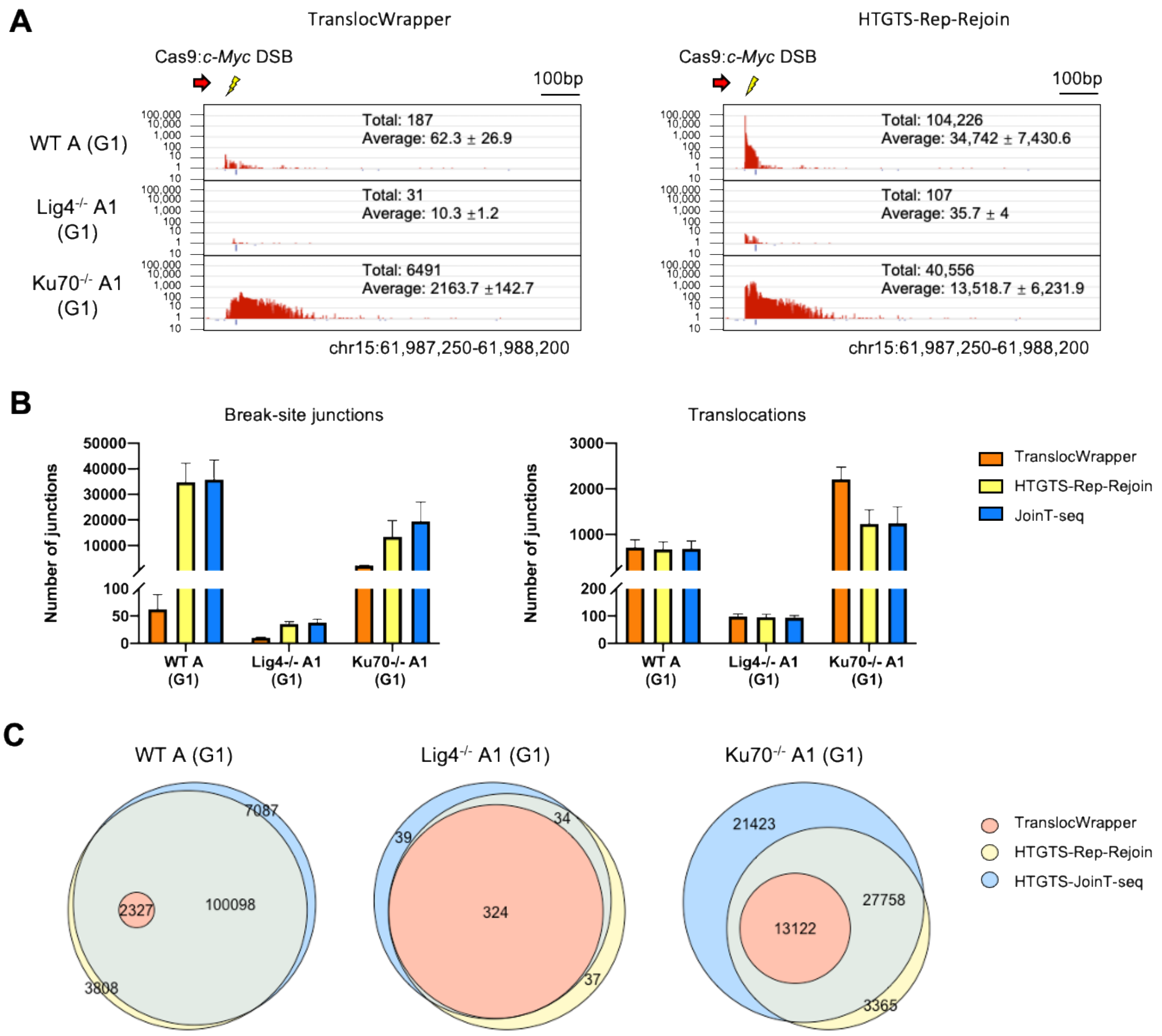


**Fig. S2. Comparison of TranslocWrapper vs. HTGTS-Rep-Rejoin and HTGTS-JoinT-seq pipelines to identify G1-phase bait DSB rejoining events.** A) *c-Myc* bait break-site profiles for WT A, Lig4$^{-/-}$A1, and Ku70$^{-/-}$A1 Abl lines analyzed by TranslocWrapper vs HTGTS-Rep-Rejoin pipelines; see Fig. 1D to compare with HTGTS-JoinT-seq. Total and Mean ±SD junctions shown ($n$=3). B) Comparison of break-site and translocation junctions from the above Abl lines analyzed by TranslocWrapper, HTGTS-Rep-Rejoin or HTGTS-JoinT-seq pipelines ($n$=3; Mean ±SD are shown). C) Venn diagram displaying total junctions identified from the above Abl lines analyzed by TranslocWrapper, HTGTS-Rep-Rejoin or HTGTS-JoinT-seq pipelines. See Methods for details on HTGTS-Rep-Rejoin.

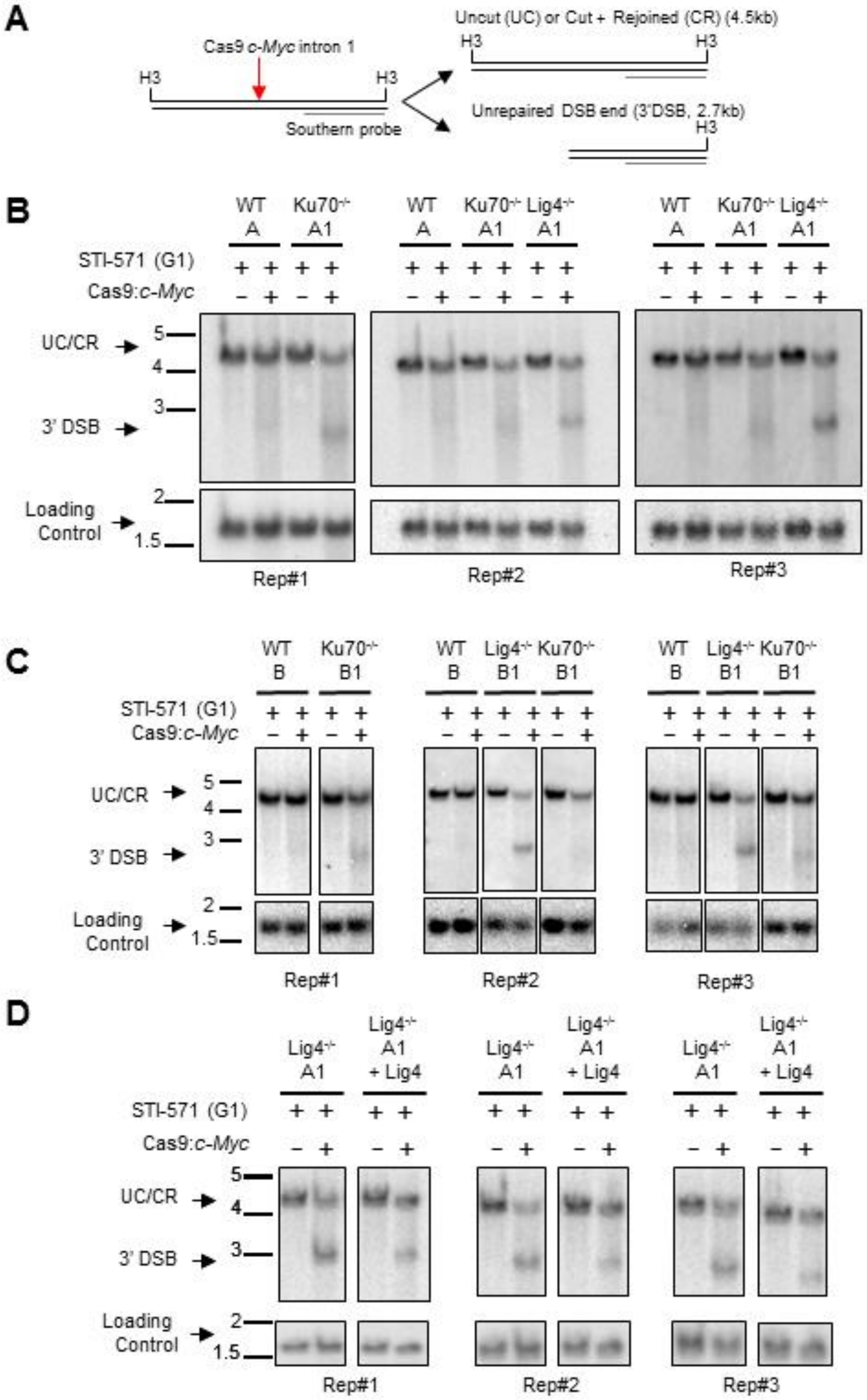


**Fig. S3. Southern blot analysis of cleavage at Cas9:*c-Myc* bait DSBs in WT, Ku70$^{-/-}$, Lig4$^{-/-}$ A and B clone set Abl cells.** A) Cas9 Cleavage at *c-Myc* locus Southern blot strategy. A probe is directed against the 3’ side of the Cas9:*c-Myc* targeting site and can detect products that represent either uncut (UC), cut + rejoined (CR), or a smaller molecular weight band corresponding to unrepaired DSBs (3’ DSB). A probe against 53bp1 served as the loading control after membrane stripping (Table S4); (H3 = HindIII). B-C) Cleavage of Cas9:*c-Myc* in WT, Ku70$^{-/-}$, Lig4$^{-/-}$ A and B Abl clone sets are assessed by Southern blot (3 separate experiments each). Corresponding HTGTS-JoinT-seq data are shown in Fig. 1. D) Cas9 bait DSB cutting detected in Lig4$^{-/-}$A1, Lig4$^{-/-}$A1 + Lig4 from three independent experiments; corresponding HTGTS-JoinT-seq data are shown in Fig. 2.

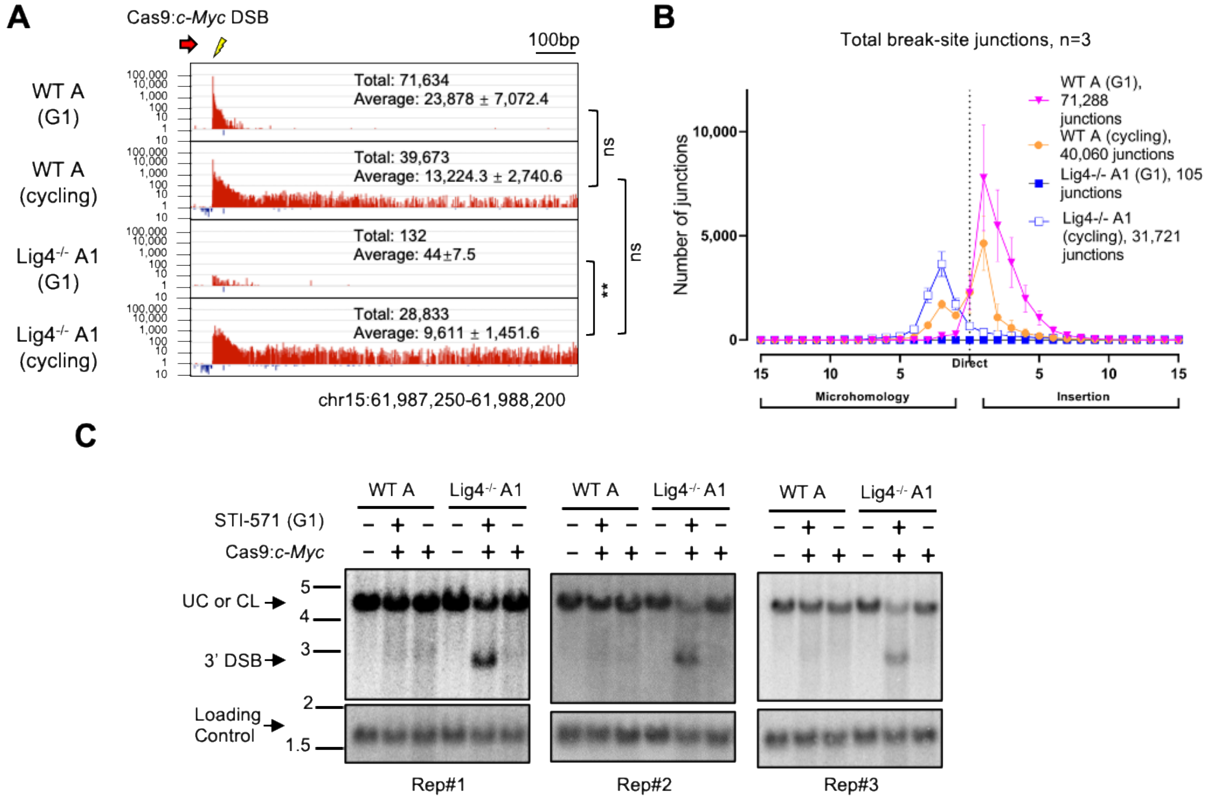


**Fig. S4. WT A Abl cells and Lig4$^{-/-}$A1 robustly rejoin the Cas9:*c-Myc* bait DSB under G1-arrest and cycling conditions.** A) Cas9:*c-Myc* bait break-site rejoining profiles for the WT A Lig4$^{-/-}$A1 Abl cells; junctions are plotted similar to Fig. 1D (*n*=3 each, One-way ANOVA + post-hoc Tukey's test, *P*<0.01, **). B) Microhomology and insertion usage of break-site junctions are plotted with indicated lengths (*n*=3; Mean ±SD). C) Cleavage of Cas9:*c-Myc* in WT A, Lig4$^{-/-}$A1 Abl lines under G1-arrest and cycling conditions are assessed by Southern blot (3 separate experiments each) as detailed in Fig S3.

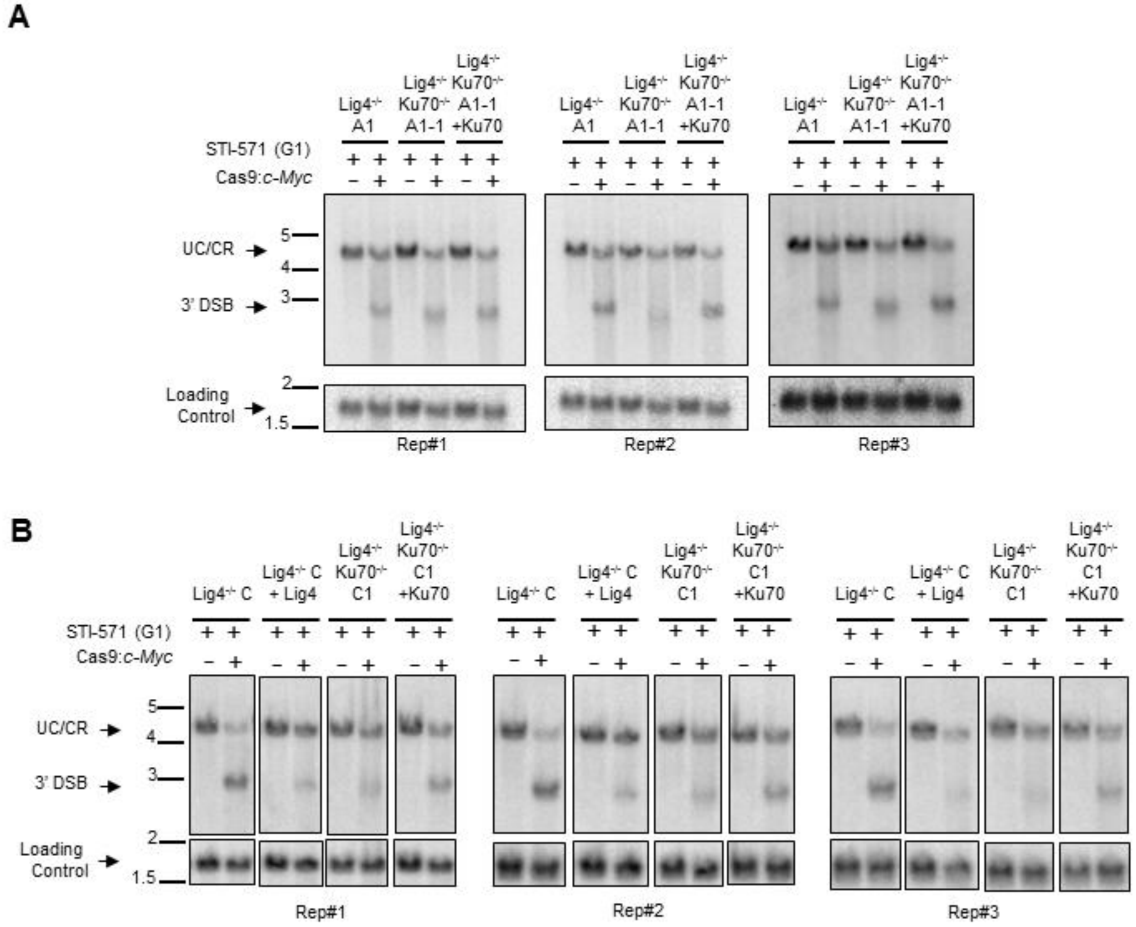


**Fig. S5. Southern blot analysis of cleavage at Cas9:*c-Myc* bait DSBs.** See Fig. S3A for the strategy. A) Cas9 bait DSB detection in Lig4$^{-/-}$A1, Lig4$^{-/-}$Ku70$^{-/-}$A1-1, or Lig4$^{-/-}$Ku70$^{-/-}$A1-1 +Ku70 Abl cells (clone A set) was assessed by Southern blot; corresponding HTGTS-JoinT-seq data are shown in Fig. 2. B) Cleavage at the Cas9 DSB site in Lig4$^{-/-}$C, Lig4$^{-/-}$C +Lig4, Lig4$^{-/-}$Ku70$^{-/-}$C1 or Lig4$^{-/-}$Ku70$^{-/-}$C1 +Ku70 Abl cells (clone C set) assessed by Southern blot; corresponding HTGTS-JoinT-seq data are shown in Fig. S6.

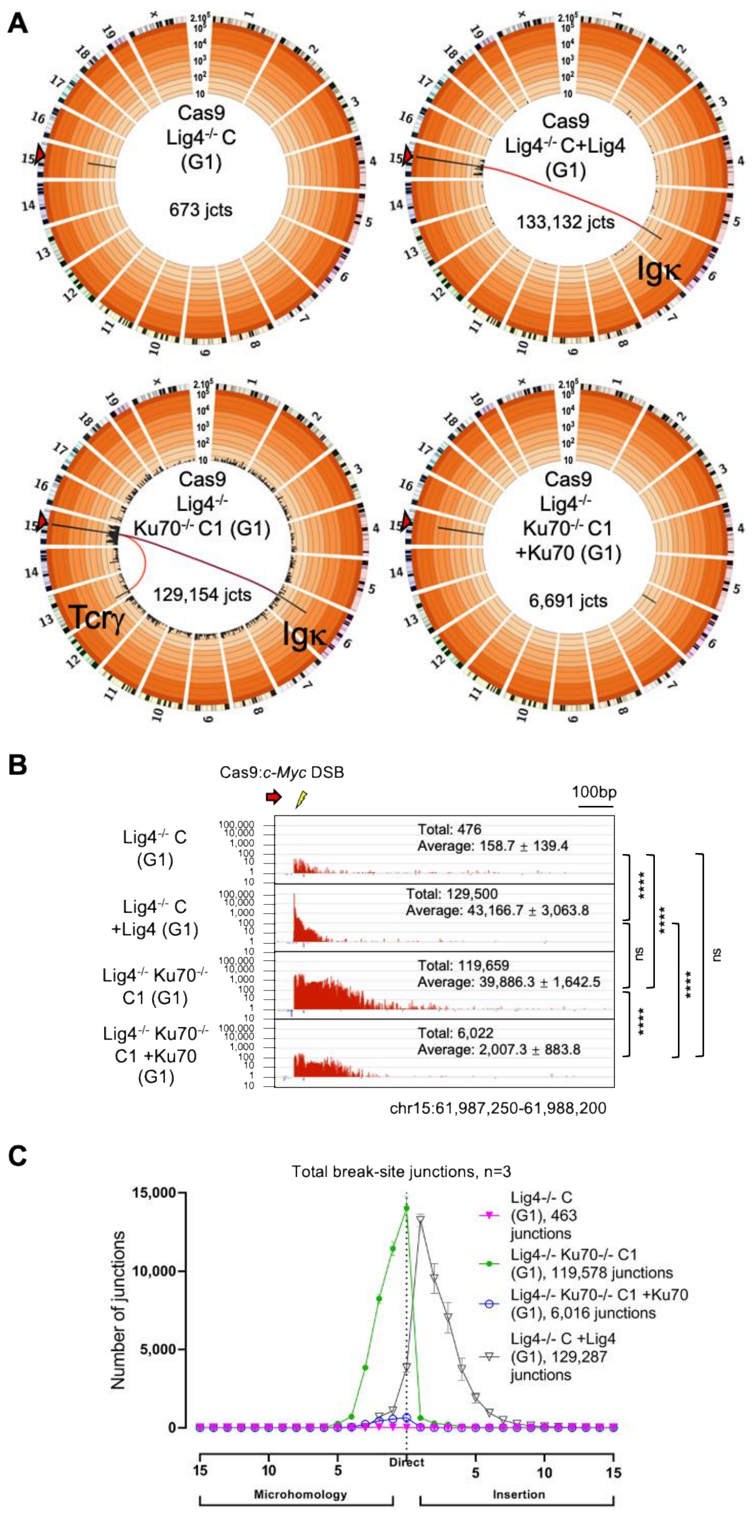


**Fig. S6. Ku70 suppresses Cas9:*c-Myc* bait DSB rejoining and translocations in G1-arrested, Lig4$^{-/-}$C, Abl cells.** A) Circos plots, as described in Fig. 1A, for in Lig4$^{-/-}$C, Lig4$^{-/-}$Ku70$^{-/-}$C1, Lig4$^{-/-}$Ku70$^{-/-}$C1 +Ku70, or Lig4$^{-/-}$C +Lig4 Abl cells. B) Cas9:*c-Myc* bait break-site rejoining profiles for the same backgrounds as in (A). Junctions plotted are similar to Fig. 2B ($n$=3 each, One-way ANOVA with post-hoc Tukey's test, $P$<0.0001, ****). C) Microhomology and insertion usage of break-site junctions are plotted with indicated lengths ($n$=3 for each clone, data represents Mean ±SD).

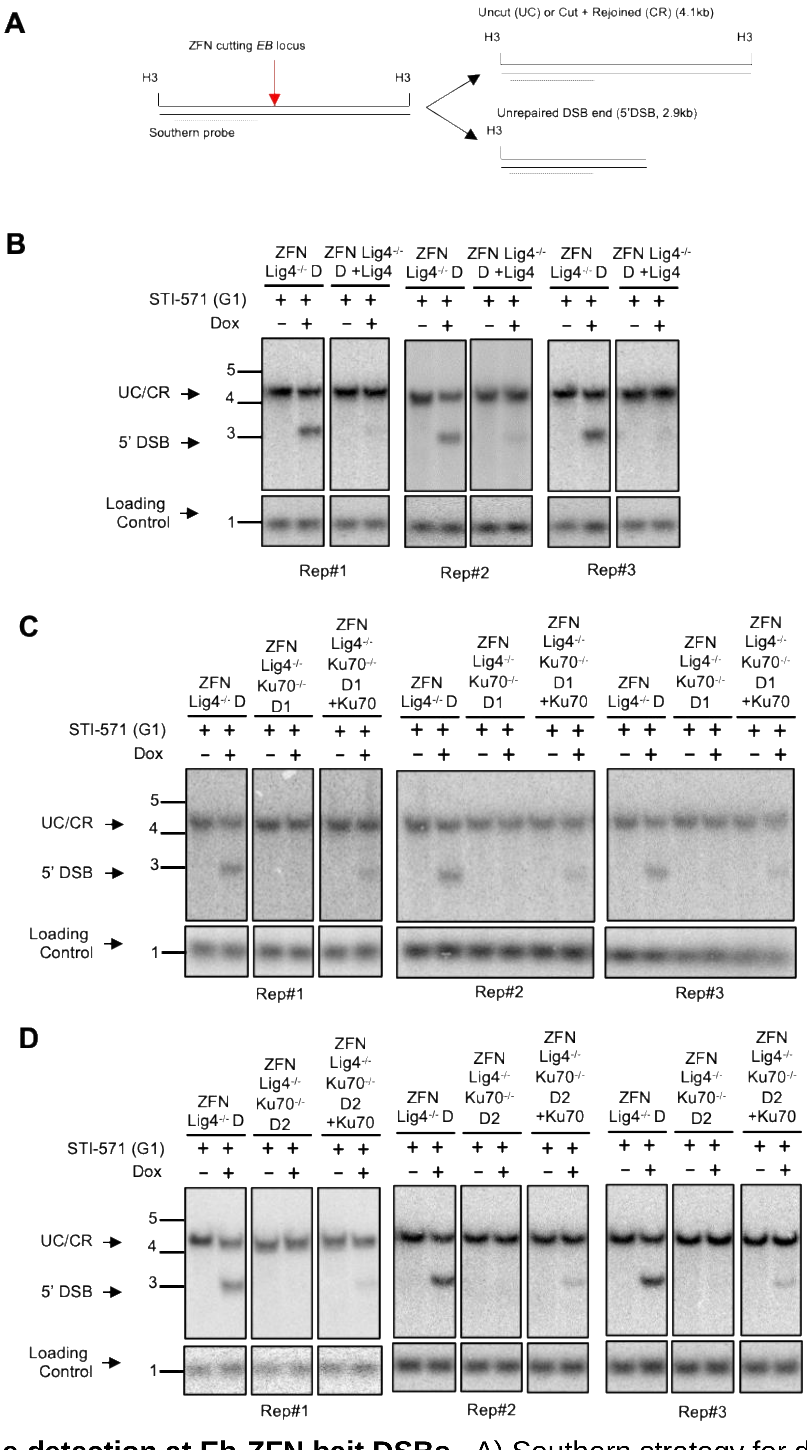


**Fig. S7. Cleavage detection at Eb-ZFN bait DSBs.** A) Southern strategy for detecting ZFN cleavage at *Eb* locus; H3 = HindIII. A 5' probe of the break-site can detect products: uncut (UC), cut and re-ligated (CL), or unrepaired DSBs (5' DSB). The ERag probe served as loading control after membrane stripping (Table S4). B-D) ZFN cleavage at *Eb* locus in Lig4$^{-/-}$D, Lig4$^{-/-}$D +Lig4, Lig4$^{-/-}$Ku70$^{-/-}$D1 and D2, or Lig4$^{-/-}$Ku70$^{-/-}$D1 and D2 +Ku70 G1-arrested Abl lines; corresponding HTGTS-JoinT-seq data are shown in Fig. 3 and Fig. S8.

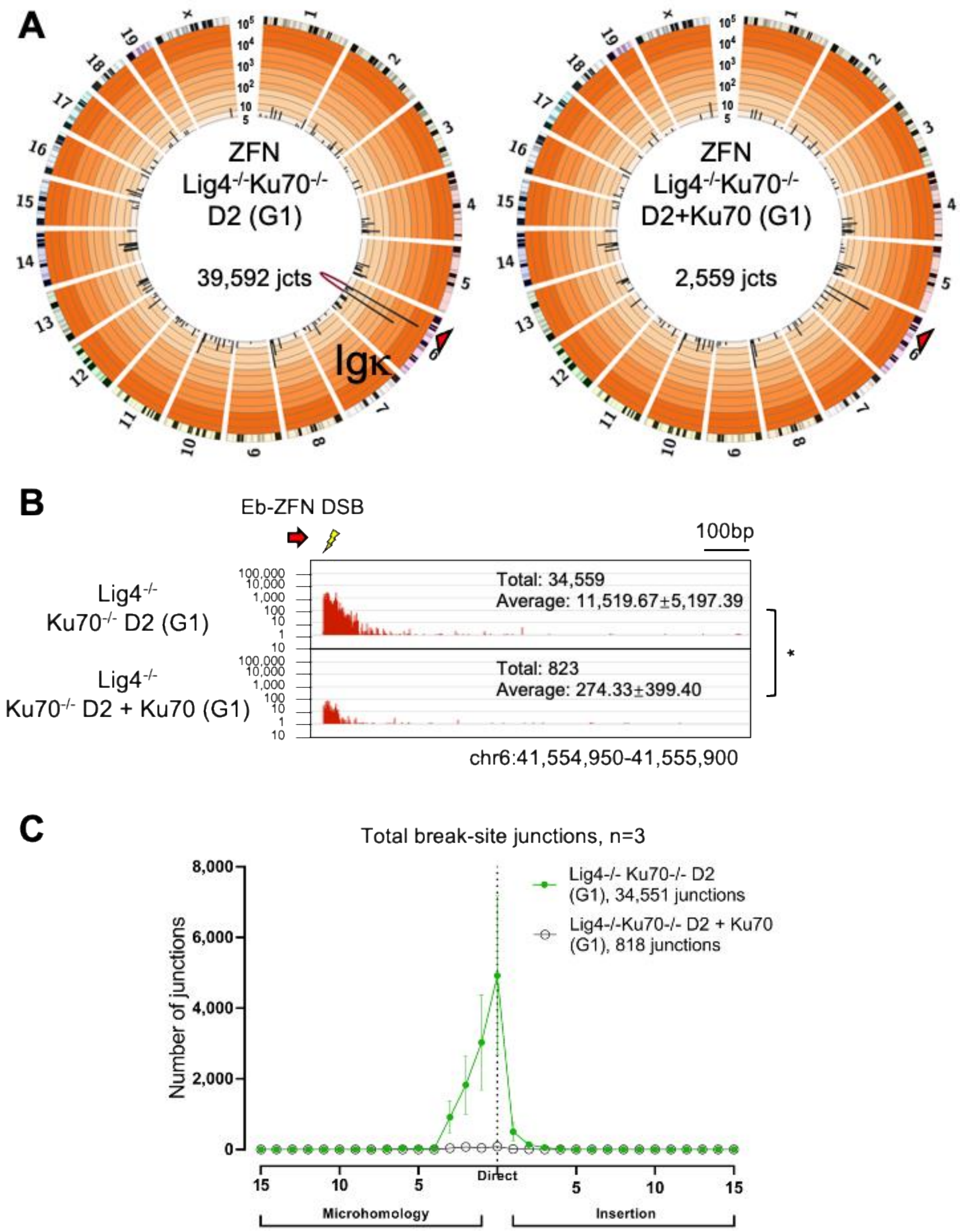


**Fig. S8. Ku70 suppresses Eb-ZFN bait DSB rejoining and translocations G1-arrested Lig4$^{-/-}$D Abl cells.** A) Circos plots as described in Fig. 3A. Deletion of Ku70 in Eb-ZFN Lig4$^{-/-}$D Abl line increases repair to DSBs genome-wide from the Eb-ZFN DSB in G1-arrested cells. B) Eb-ZFN bait break-site rejoining profiles from Lig4$^{-/-}$Ku70$^{-/-}$D2, or Lig4$^{-/-}$Ku70$^{-/-}$D2 +Ku70 Abl cells. Junctions are plotted similar to Fig. 3B (*n*=3 for each clone, unpaired t-test, *P*<0.05, *). C) Microhomology and insertion usage of break-site junctions are plotted with indicated lengths (*n*=3 for each clone, data represents Mean ±SD).

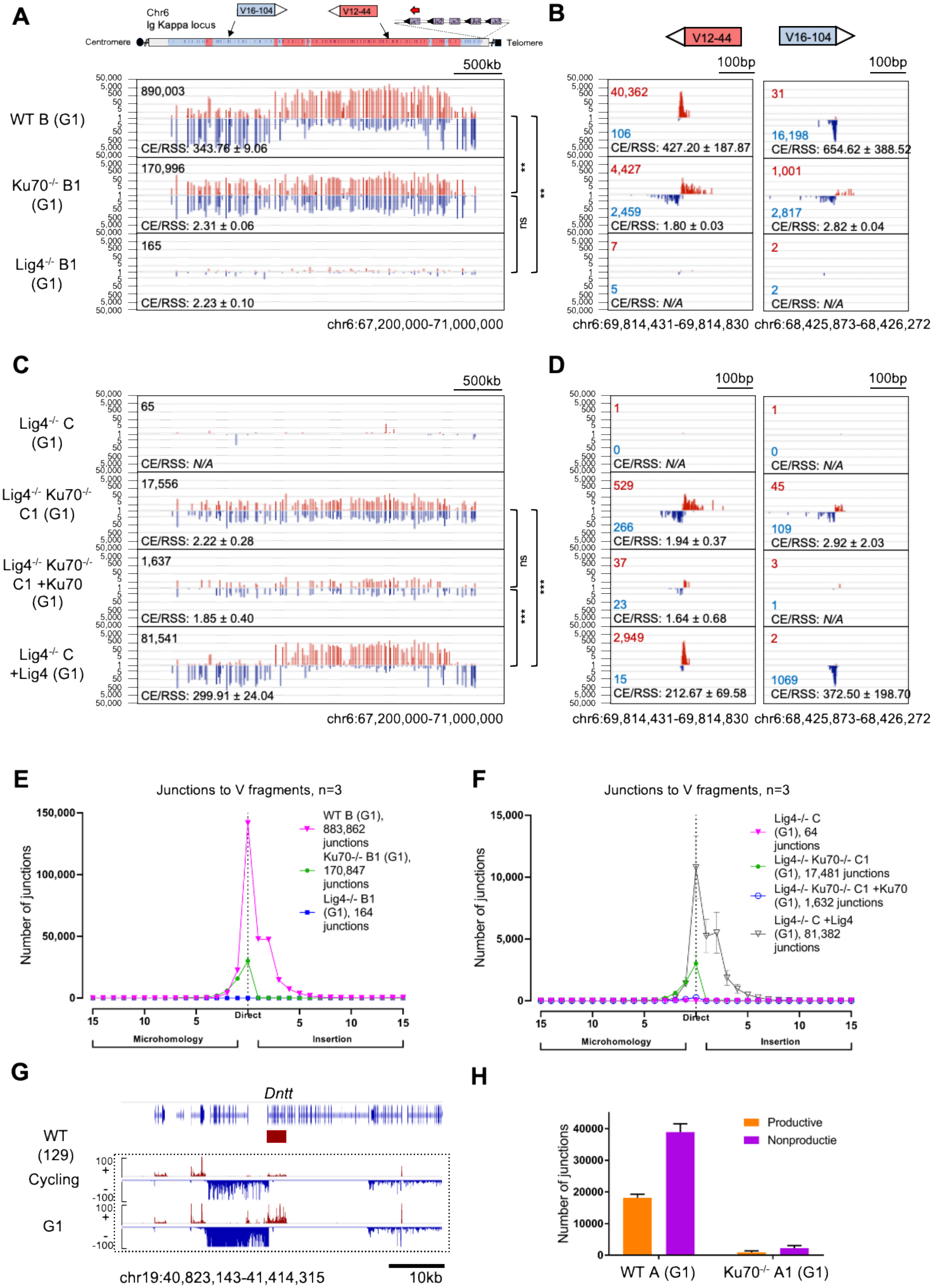
A
Chr6
Ig Kappa locus
V16-104
V12-44
Centromere
Telomere
500kb
WT B (G1)
890,003
CE/RSS: 343.76 ± 9.06
Ku70-/- B1 (G1)
170,996
CE/RSS: 2.31 ± 0.06
Lig4-/- B1 (G1)
165
CE/RSS: 2.23 ± 0.10
**
**
ns
chr6:67,200,000-71,000,000
B
V12-44
V16-104
100bp
40,362
106
CE/RSS: 427.20 ± 187.87
31
16,198
CE/RSS: 654.62 ± 388.52
4,427
2,459
CE/RSS: 1.80 ± 0.03
1,001
2,817
CE/RSS: 2.82 ± 0.04
7
5
CE/RSS: N/A
2
2
CE/RSS: N/A
chr6:69,814,431-69,814,830
chr6:68,425,873-68,426,272
C
Lig4-/- C (G1)
65
CE/RSS: N/A
Lig4-/- Ku70-/- C1 (G1)
17,556
CE/RSS: 2.22 ± 0.28
Lig4-/- Ku70-/- C1 +Ku70 (G1)
1,637
CE/RSS: 1.85 ± 0.40
Lig4-/- C +Lig4 (G1)
81,541
CE/RSS: 299.91 ± 24.04
ns
***
***
chr6:67,200,000-71,000,000
D
1
0
CE/RSS: N/A
1
0
CE/RSS: N/A
529
266
CE/RSS: 1.94 ± 0.37
45
109
CE/RSS: 2.92 ± 2.03
37
23
CE/RSS: 1.64 ± 0.68
3
1
CE/RSS: N/A
2,949
15
CE/RSS: 212.67 ± 69.58
2
1069
CE/RSS: 372.50 ± 198.70
chr6:69,814,431-69,814,830
chr6:68,425,873-68,426,272
E
Junctions to V fragments, n=3
Number of junctions
150,000
100,000
50,000
0
WT B (G1), 883,862 junctions
Ku70-/- B1 (G1), 170,847 junctions
Lig4-/- B1 (G1), 164 junctions
Direct
Microhomology
Insertion
F
Junctions to V fragments, n=3
Number of junctions
15,000
10,000
5,000
0
Lig4-/- C (G1), 64 junctions
Lig4-/- Ku70-/- C1 (G1), 17,481 junctions
Lig4-/- Ku70-/- C1 +Ku70 (G1), 1,632 junctions
Lig4-/- C +Lig4 (G1), 81,382 junctions
Direct
Microhomology
Insertion
G
Dntt
WT (129)
Cycling
G1
100
-100
chr19:40,823,143-41,414,315
10kb
H
Number of junctions
40000
30000
20000
10000
0
Productive
Nonproductie
WT A (G1)
Ku70-/- A1 (G1)

**Fig. S9. Ku suppresses A-EJ of RAG DSBs to each other; otherwise, A-EJ employs a translocation-based mechanism for their repair.** A,C) Jκ1CE bait HTGTS-V(D)J-seq comparing WT B, Ku70$^{-/-}$B1, and Lig4$^{-/-}$B1 Abl cells (A) or Lig4$^{-/-}$C, Lig4$^{-/-}$C +Lig4, Lig4$^{-/-}$Ku70$^{-/-}$C1, Lig4$^{-/-}$Ku70$^{-/-}$C1 +Ku70 Abl cells (C) (See Figure 4A legend; One-way ANOVA to compare the ratios CE/RSS with post-hoc Tukey's test, *P<0.01,* **, *P*<0.001, ***). B,D) Zoom-in of select V gene segments from (A,C). E-F) Microhomology and insertion usage in junctions joining from Jκ1 to Vκ segments are plotted with indicated lengths (*n*=3 for each clone, data represents Mean ±SD). G) Transcriptional activity (GRO-seq) of Tdt (DNTT) in cycling and G1-arrested Abl cells. H) Productive and nonproductive junctions from WT A and Ku70$^{-/-}$A1 Abl Jκ1 CE bait to Vκ CEs.

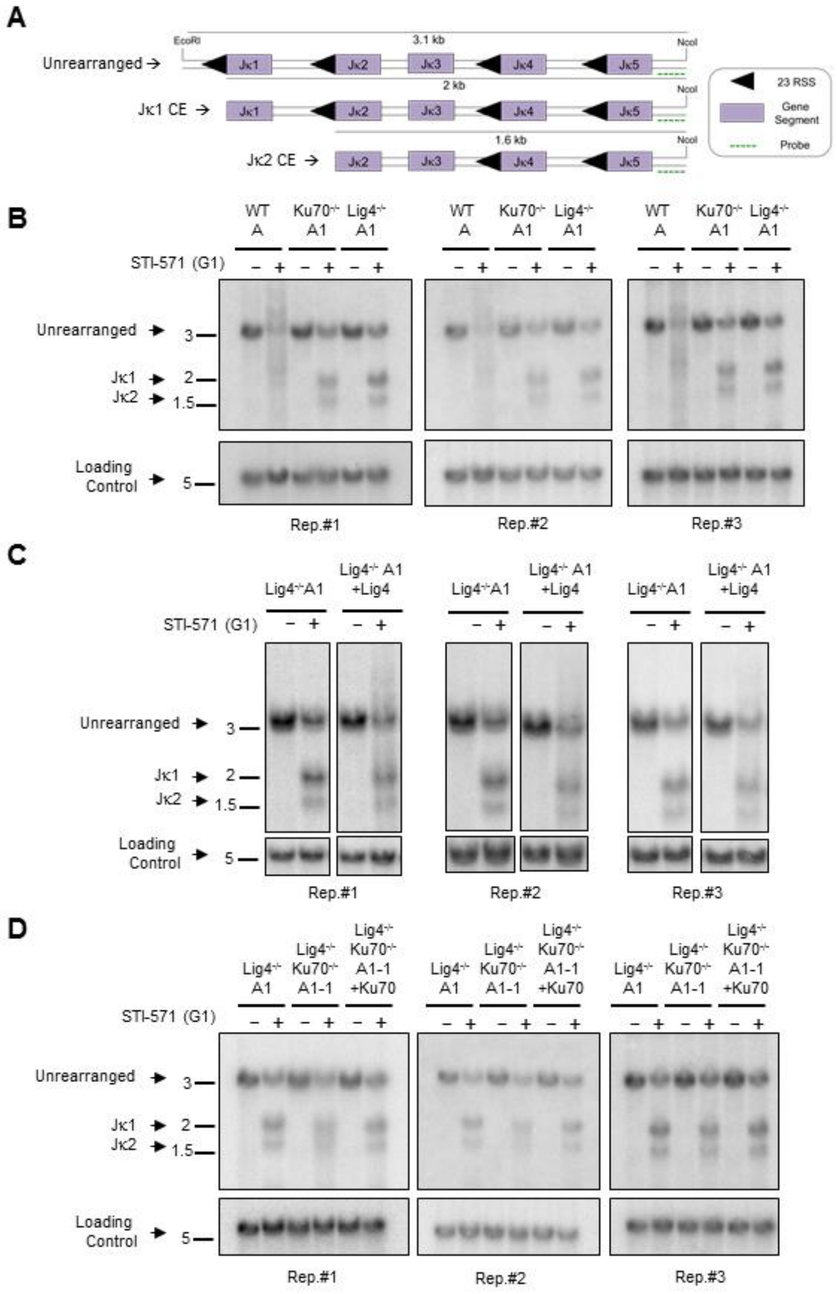


**Fig. S10. Southern blot analysis of Jκ gene segment cleavage by RAG.** A) A 3' end probe downstream of Jκ5 to monitor RAG cleavage; note that Jκ3 does not cut. B) The smear extending above and below the germline band in WT Abl cells are heterogeneous Vκ-Jκ recombination products, whereas accumulation of unrepaired Jκ coding ends are shown for Ku70$^{-/-}$ and Lig4$^{-/-}$ Abl cells. ERag probe was used as a loading control. C-D) RAG cleavage of Jκ gene segments in the WT A, Ku70$^{-/-}$A1, Lig4$^{-/-}$A1, Lig4$^{-/-}$A1 +Lig4, Lig4$^{-/-}$Ku70$^{-/-}$A1-1, and Lig4$^{-/-}$Ku70$^{-/-}$A1-1 +Ku70 Abl cells from three independent replications. Related HTGTS-V(D)J-seq data are shown in Fig. 4.

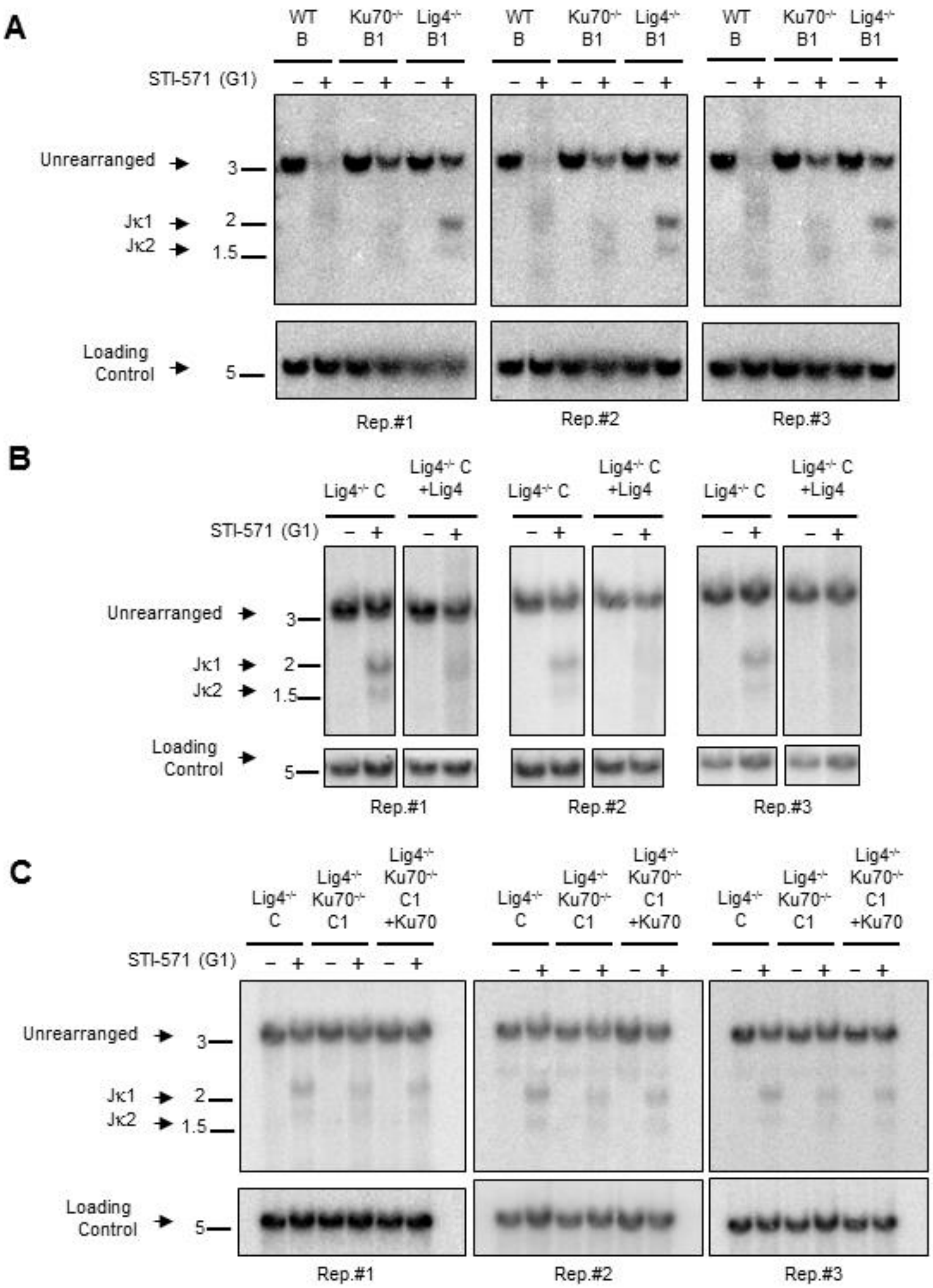


**Fig. S11. Jκ gene segment cleavage by RAG.** A) RAG cleavage of Jκ gene segments in WT B, Ku70$^{-/-}$B1, Lig4$^{-/-}$B1 Abl cells by Southern blot. Related LAM-HTGTS data are shown in Fig. S10A. B-C) RAG cleavage of Jκ gene segments assayed in Lig4$^{-/-}$C, Lig4$^{-/-}$C +Lig4, Lig4$^{-/-}$Ku70$^{-/-}$C1, and Lig4$^{-/-}$Ku70$^{-/-}$C1 +Ku70 Abl cells by Southern blot. Three independent experiments shown. Related HTGTS-V(D)J-seq data are shown in Fig. S10C.

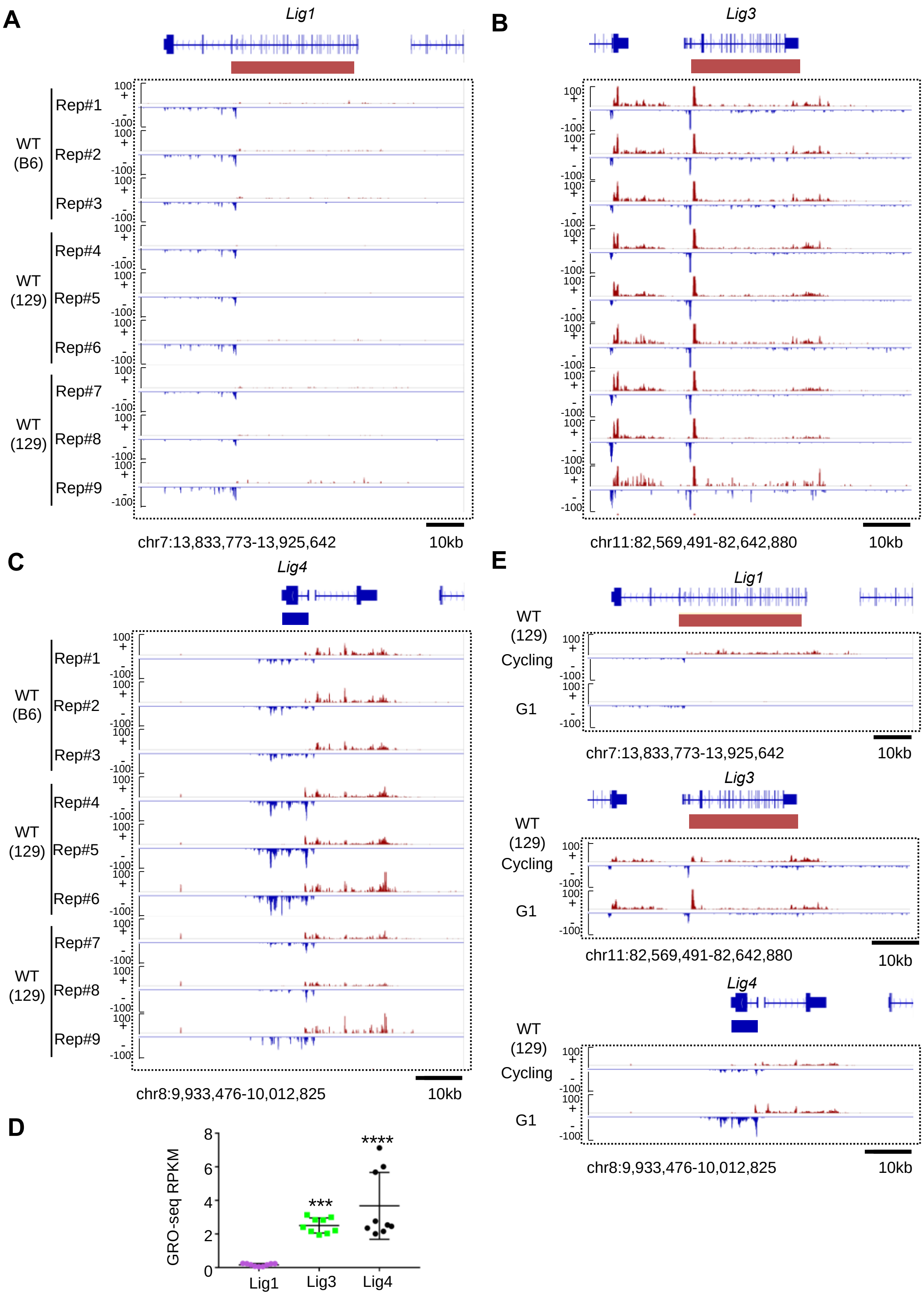


**Fig S12. Transcription activity of Lig1, Lig3 and Lig4 in G1-arrestd or cycling Abl cells.** A-C) Transcription activity of Lig1, Lig3 and Lig4 from independent GRO-seq experiments and mouse strain (B6, 129) indicated. (Rep#1-3 are using GSM4593316-GSM4593318 from Dai et al., 2021; Rep#4-6 are using GSM4240123, GSM4252097, GSM4252099 from Ba et al., 2020; Rep#7-9 are using GSM4252101, GSM4252103, GSM4252105 from Ba et al., 2020) D) Transcription activity measured by reads per kilobase per million (RPKM) from GRO-seq experiments is plotted for Lig1, Lig3 and Lig4. (One-way ANOVA with post-hoc Tukey's test, Lig3 vs. Lig1, $P < 0.001$, ***, Lig4 vs. Lig1, $P < 0.0001$, ****; $n$=9, data represents Mean±SD). E) Transcription activity of Lig1, Lig3 and Lig4 from cycling or G1-arrested Abl cells.

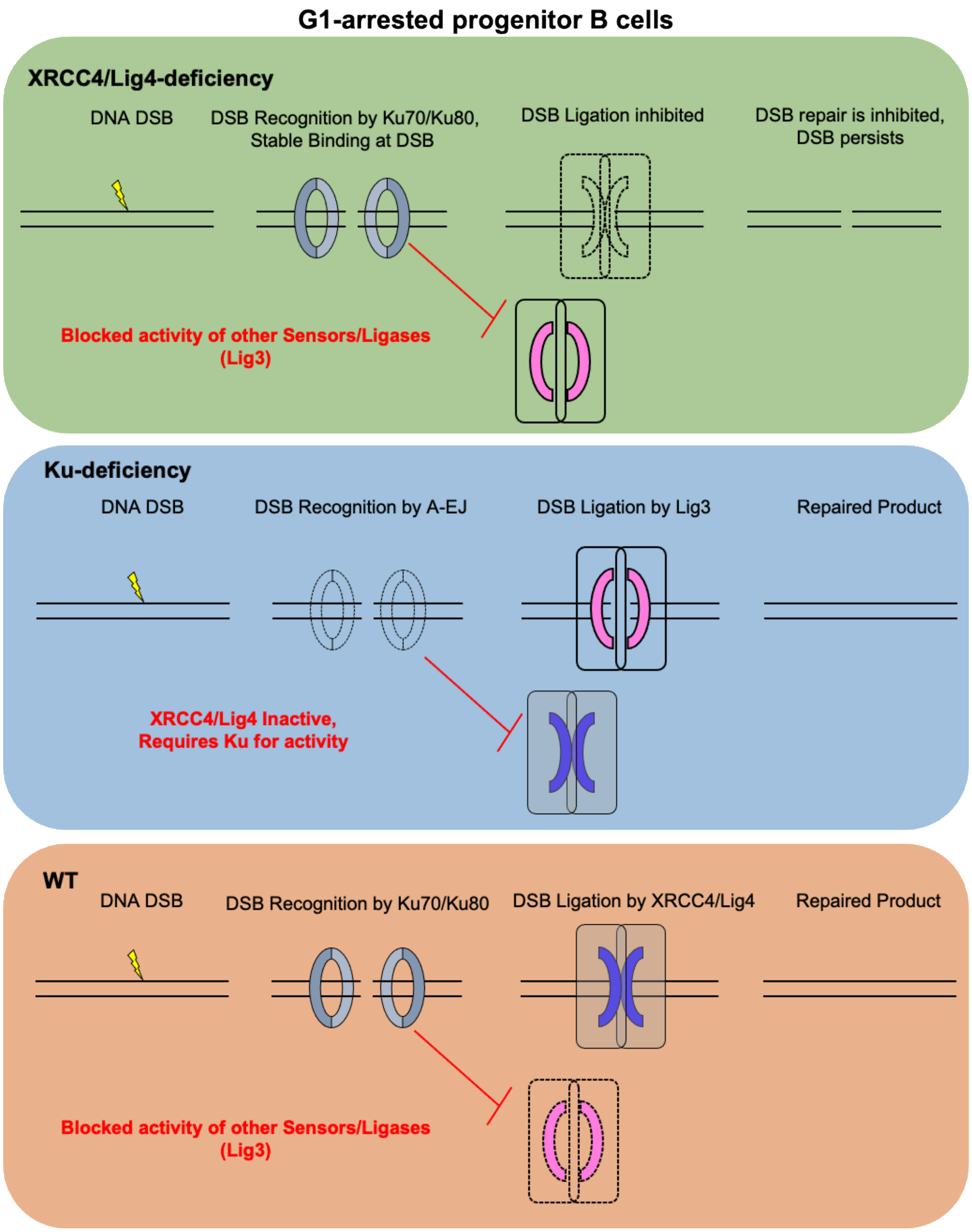


**Fig S13. Underlying mechanism of Ku protein suppressing Lig3-mediated Alternative End-Joining in G1-arrested pro-B cells.** We propose that the Ku complex senses any DSB generated in G1-arrested Abl cells, regardless of Lig4 status. If both the Ku complex and XRCC4/Ligase4 complex are present, then repair of DSBs occurs faithfully and exclusively through C-NHEJ. However, in absence of Ligase4, Ku would bind these ends and prevent access to Ligase3 (Lig3) activity from completing their repair, thereby leading to a complete block in both C-NHEJ and A-EJ in G1 where Ligase1 expression is low.

### Table S1. Cas9:*c-Myc* and Eb-ZFN HTGTS-JoinT-seq libraries information

| Genotype | Breaks | G1 / cycling | Rep | Reads | Total Junctions * | Break-site junctions ** | Translocations *** | % of Breaksite junctions | Igκ Translocations **** |
|---|---|---|---|---|---|---|---|---|---|
| **Figure 1A,D** | | | | | | | | | |
| WT A | Cas9:c-Myc | G1 | 1 | 396,387 | 30,164 | 29,383 | 781 | 97.4% | 124 |
| | Cas9:c-Myc | G1 | 2 | 396,387 | 44,977 | 44,190 | 787 | 98.3% | 103 |
| | Cas9:c-Myc | G1 | 3 | 396,387 | 34,371 | 33,874 | 497 | 98.6% | 62 |
| Lig4-/- A1 | Cas9:c-Myc | G1 | 1 | 396,387 | 146 | 43 | 103 | 29.5% | 0 |
| | Cas9:c-Myc | G1 | 2 | 396,387 | 124 | 32 | 92 | 25.8% | 2 |
| | Cas9:c-Myc | G1 | 3 | 396,387 | 127 | 40 | 87 | 31.5% | 1 |
| Ku70-/- A1 | Cas9:c-Myc | G1 | 1 | 396,387 | 23,198 | 22,265 | 933 | 96.0% | 105 |
| | Cas9:c-Myc | G1 | 2 | 396,387 | 12,668 | 11,028 | 1,640 | 87.1% | 277 |
| | Cas9:c-Myc | G1 | 3 | 396,387 | 26,437 | 25,273 | 1,164 | 95.6% | 90 |
| **Figure 1B,E** | | | | | | | | | |
| WT B | Cas9:c-Myc | G1 | 1 | 600,000 | 28,028 | 26,555 | 1,473 | 94.7% | 284 |
| | Cas9:c-Myc | G1 | 2 | 600,000 | 29,640 | 28,266 | 1,374 | 95.4% | 282 |
| | Cas9:c-Myc | G1 | 3 | 600,000 | 55,809 | 53,214 | 2,595 | 95.4% | 344 |
| Lig4-/- B1 | Cas9:c-Myc | G1 | 1 | 600,000 | 155 | 60 | 95 | 38.7% | 1 |
| | Cas9:c-Myc | G1 | 2 | 600,000 | 140 | 39 | 101 | 27.9% | 3 |
| | Cas9:c-Myc | G1 | 3 | 600,000 | 128 | 23 | 105 | 18.0% | 1 |
| Ku70-/- B1 | Cas9:c-Myc | G1 | 1 | 600,000 | 47,748 | 42,968 | 4,780 | 90.0% | 1,577 |
| | Cas9:c-Myc | G1 | 2 | 600,000 | 25,950 | 23,440 | 2,510 | 90.3% | 1,023 |
| | Cas9:c-Myc | G1 | 3 | 600,000 | 51,862 | 46,020 | 5,842 | 88.7% | 1,913 |
| **Figure 2A,B** | | | | | | | | | |
| Lig4-/- A1 | Cas9:c-Myc | G1 | 1 | 201,858 | 86 | 50 | 36 | 58.1% | 1 |
| | Cas9:c-Myc | G1 | 2 | 201,858 | 411 | 358 | 53 | 87.1% | 1 |
| | Cas9:c-Myc | G1 | 3 | 201,858 | 82 | 16 | 66 | 19.5% | 2 |
| Lig4-/- A1 +Lig4 | Cas9:c-Myc | G1 | 1 | 201,858 | 8,642 | 8,106 | 536 | 93.8% | 133 |
| | Cas9:c-Myc | G1 | 2 | 201,858 | 12,595 | 11,780 | 815 | 93.5% | 206 |
| | Cas9:c-Myc | G1 | 3 | 201,858 | 11,456 | 10,584 | 872 | 92.4% | 281 |
| Lig4-/- Ku70-/- A1-1 | Cas9:c-Myc | G1 | 1 | 201,858 | 9,082 | 8,515 | 567 | 93.8% | 38 |
| | Cas9:c-Myc | G1 | 2 | 201,858 | 6,412 | 5,974 | 438 | 93.2% | 39 |
| | Cas9:c-Myc | G1 | 3 | 201,858 | 13,431 | 11,510 | 1,921 | 85.7% | 246 |
| Lig4-/- Ku70-/- A1-1 +Ku70 | Cas9:c-Myc | G1 | 1 | 201,858 | 106 | 75 | 31 | 70.8% | 1 |
| | Cas9:c-Myc | G1 | 2 | 201,858 | 128 | 101 | 27 | 78.9% | 6 |
| | Cas9:c-Myc | G1 | 3 | 201,858 | 278 | 157 | 121 | 56.5% | 14 |
| **Figure 3A,B** | | | | | | | | | |
| Lig4-/- D | Eb-ZFN | G1 | 1 | 600,000 | 339 | 279 | 60 | 82.3% | 6 |
| | Eb-ZFN | G1 | 2 | 600,000 | 660 | 566 | 94 | 85.8% | 7 |
| | Eb-ZFN | G1 | 3 | 600,000 | 665 | 607 | 58 | 91.3% | 8 |
| Lig4-/- D +Lig4 | Eb-ZFN | G1 | 1 | 600,000 | 9,709 | 9,497 | 212 | 97.8% | 137 |
| | Eb-ZFN | G1 | 2 | 600,000 | 9,102 | 8,898 | 204 | 97.8% | 130 |
| | Eb-ZFN | G1 | 3 | 600,000 | 8,865 | 8,655 | 210 | 97.6% | 151 |
| Lig4-/- Ku70-/- D1 | Eb-ZFN | G1 | 1 | 600,000 | 17,254 | 15,925 | 1,329 | 92.3% | 979 |
| | Eb-ZFN | G1 | 2 | 600,000 | 16,870 | 15,368 | 1,502 | 91.1% | 1,126 |
| | Eb-ZFN | G1 | 3 | 600,000 | 17,245 | 15,698 | 1,547 | 91.0% | 1,165 |
| Lig4-/- Ku70-/- D1 +Ku70 | Eb-ZFN | G1 | 1 | 600,000 | 1,207 | 906 | 301 | 75.1% | 53 |
| | Eb-ZFN | G1 | 2 | 600,000 | 988 | 686 | 302 | 69.4% | 41 |
| | Eb-ZFN | G1 | 3 | 600,000 | 839 | 567 | 272 | 67.6% | 35 |

| | | | | | | | | | |
|---|---|---|---|---|---|---|---|---|---|
| **Figure S4A** | | | | | | | | | |
| WT A | Cas9:c-Myc | G1 | 1 | 310,675 | 15,888 | 15,748 | 140 | 99.1% | 7 |
| | Cas9:c-Myc | G1 | 2 | 310,675 | 27,583 | 27,275 | 308 | 98.9% | 38 |
| | Cas9:c-Myc | G1 | 3 | 310,675 | 28,933 | 28,611 | 322 | 98.9% | 34 |
| WT A | Cas9:c-Myc | cycling | 1 | 310,675 | 14,024 | 13,755 | 269 | 98.1% | 0 |
| | Cas9:c-Myc | cycling | 2 | 310,675 | 10,909 | 10,696 | 213 | 98.0% | 0 |
| | Cas9:c-Myc | cycling | 3 | 310,675 | 16,293 | 16,026 | 267 | 98.4% | 2 |
| Lig4-/- A1 | Cas9:c-Myc | G1 | 1 | 310,675 | 74 | 43 | 31 | 58.1% | 0 |
| | Cas9:c-Myc | G1 | 2 | 310,675 | 100 | 52 | 48 | 52.0% | 11 |
| | Cas9:c-Myc | G1 | 3 | 310,675 | 85 | 37 | 48 | 43.5% | 0 |
| Lig4-/- A1 | Cas9:c-Myc | cycling | 1 | 310,675 | 10,022 | 9,552 | 470 | 95.3% | 0 |
| | Cas9:c-Myc | cycling | 2 | 310,675 | 13,060 | 12,534 | 526 | 96.0% | 1 |
| | Cas9:c-Myc | cycling | 3 | 310,675 | 10,829 | 10,311 | 518 | 95.2% | 0 |
| **Figure S6A,B** | | | | | | | | | |
| Lig4-/- C | Cas9:c-Myc | G1 | 1 | 273,198 | 138 | 59 | 79 | 42.8% | 2 |
| | Cas9:c-Myc | G1 | 2 | 273,198 | 385 | 318 | 67 | 82.6% | 0 |
| | Cas9:c-Myc | G1 | 3 | 273,198 | 150 | 99 | 51 | 66.0% | 0 |
| Lig4-/- C +Lig4 | Cas9:c-Myc | G1 | 1 | 273,198 | 47,225 | 46,297 | 928 | 98.0% | 68 |
| | Cas9:c-Myc | G1 | 2 | 273,198 | 41,481 | 40,174 | 1,307 | 96.8% | 79 |
| | Cas9:c-Myc | G1 | 3 | 273,198 | 44,440 | 43,029 | 1,411 | 96.8% | 78 |
| Lig4-/- Ku70-/- C1 | Cas9:c-Myc | G1 | 1 | 273,198 | 42,340 | 39,082 | 3,258 | 92.3% | 704 |
| | Cas9:c-Myc | G1 | 2 | 273,198 | 44,798 | 41,776 | 3,022 | 93.3% | 594 |
| | Cas9:c-Myc | G1 | 3 | 273,198 | 42,026 | 38,801 | 3,225 | 92.3% | 786 |
| Lig4-/- Ku70-/- C1 +Ku70 | Cas9:c-Myc | G1 | 1 | 273,198 | 2,204 | 1,942 | 262 | 88.1% | 35 |
| | Cas9:c-Myc | G1 | 2 | 273,198 | 3,167 | 2,922 | 245 | 92.3% | 30 |
| | Cas9:c-Myc | G1 | 3 | 273,198 | 1,320 | 1,158 | 162 | 87.7% | 23 |
| **Figure S8A,B** | | | | | | | | | |
| Lig4-/- Ku70-/- D2 | Eb-ZFN | G1 | 1 | 600,000 | 17,177 | 14,311 | 2,866 | 83.3% | 1,274 |
| | Eb-ZFN | G1 | 2 | 600,000 | 16,257 | 14,725 | 1,532 | 90.6% | 1,206 |
| | Eb-ZFN | G1 | 3 | 600,000 | 6,164 | 5,523 | 641 | 89.6% | 434 |
| Lig4-/- Ku70-/- D2 +Ku70 | Eb-ZFN | G1 | 1 | 600,000 | 2,202 | 735 | 1,467 | 33.4% | 37 |
| | Eb-ZFN | G1 | 2 | 600,000 | 196 | 63 | 133 | 32.1% | 2 |
| | Eb-ZFN | G1 | 3 | 600,000 | 161 | 25 | 136 | 15.5% | 2 |

* Total junctions include junctions in chr1-19, chrX, chrY, chr M

** Break-site junctions are defined for G1 cells as junctions in 950bp interval downstream of the break-site (chr15:61,987,250-61,988,200 for *c-Myc* locus, chr6:41,554,950-41,555,900 for Eb locus) and for cycling cells as junctions in a 10kb interval (chr15:61,987,250-61,997,250 for Myc locus, chr6:41,554,950-41,564,950 for Eb locus)

*** Translocations are defined as junctions outside of the break-site window

**** Igκ translocations are defined as junctions in the interval chr6:67,200,000-71,000,000

### Table S2. Resection quantification for Cas9:*c-Myc* and Eb-ZFN HTGTS-JoinT-seq libraries

| Genotype | Breaks | G1 / cycling | Rep | Rejoining junctions * | Resection mean (bp) | Resection median (bp) | Median average | Median sd | 90%** (bp) | 99%** (bp) | 99.9%** (bp) |
|---|---|---|---|---|---|---|---|---|---|---|---|
| **Figure 1D** | | | | | | | | | | | |
| WT A | Cas9:c-Myc | G1 | 1 | 29,380 | 0.5 | 0 | 0.0 | 0.0 | 0 | 12 | 46 |
| | Cas9:c-Myc | G1 | 2 | 44,189 | 0.5 | 0 | | | 0 | 12 | 41 |
| | Cas9:c-Myc | G1 | 3 | 33,871 | 0.4 | 0 | | | 0 | 10 | 35 |
| Lig4-/- A1 | Cas9:c-Myc | G1 | 1 | 41 | 36.9 | 35 | 27.5 | 6.9 | 56 | 213 | 213 |
| | Cas9:c-Myc | G1 | 2 | 31 | 30.3 | 26 | | | 52 | 135 | 135 |
| | Cas9:c-Myc | G1 | 3 | 38 | 23.7 | 21.5 | | | 52 | 122 | 122 |
| Ku70-/- A1 | Cas9:c-Myc | G1 | 1 | 22,264 | 45.3 | 25 | 28.7 | 8.1 | 111 | 173 | 291 |
| | Cas9:c-Myc | G1 | 2 | 11,025 | 52.7 | 38 | | | 111 | 169 | 276 |
| | Cas9:c-Myc | G1 | 3 | 25,266 | 38.5 | 23 | | | 104 | 165 | 306 |
| **Figure 1E** | | | | | | | | | | | |
| WT B | Cas9:c-Myc | G1 | 1 | 26,549 | 1.0 | 0 | 0.0 | 0.0 | 1 | 12 | 147 |
| | Cas9:c-Myc | G1 | 2 | 28,254 | 0.9 | 0 | | | 1 | 11 | 177 |
| | Cas9:c-Myc | G1 | 3 | 53,205 | 0.9 | 0 | | | 1 | 12 | 143 |
| Lig4-/- B1 | Cas9:c-Myc | G1 | 1 | 59 | 95.1 | 58 | 59.8 | 30.3 | 220 | 564 | 564 |
| | Cas9:c-Myc | G1 | 2 | 37 | 101.6 | 91 | | | 254 | 360 | 360 |
| | Cas9:c-Myc | G1 | 3 | 22 | 61.0 | 30.5 | | | 192 | 310 | 310 |
| Ku70-/- B1 | Cas9:c-Myc | G1 | 1 | 42,957 | 78.5 | 80 | 78.7 | 1.2 | 128 | 190 | 392 |
| | Cas9:c-Myc | G1 | 2 | 23,435 | 75.5 | 78 | | | 124 | 190 | 401 |
| | Cas9:c-Myc | G1 | 3 | 46,005 | 76.3 | 78 | | | 128 | 190 | 408 |
| **Figure 2B** | | | | | | | | | | | |
| Lig4-/- A1 | Cas9:c-Myc | G1 | 1 | 48 | 38.2 | 24 | 14.7 | 12.9 | 104 | 196 | 196 |
| | Cas9:c-Myc | G1 | 2 | 356 | 21.4 | 0 | | | 28 | 677 | 767 |
| | Cas9:c-Myc | G1 | 3 | 15 | 38.3 | 20 | | | 104 | 112 | 112 |
| Lig4-/- A1 +Lig4 | Cas9:c-Myc | G1 | 1 | 8,104 | 1.5 | 0 | 0.0 | 0.0 | 3 | 27 | 70 |
| | Cas9:c-Myc | G1 | 2 | 11,778 | 1.8 | 0 | | | 3 | 37 | 72 |
| | Cas9:c-Myc | G1 | 3 | 10,583 | 2.0 | 0 | | | 4 | 33 | 65 |
| Lig4-/- Ku70-/- A1-1 | Cas9:c-Myc | G1 | 1 | 8,515 | 33.0 | 21 | 22.3 | 3.2 | 90 | 157 | 316 |
| | Cas9:c-Myc | G1 | 2 | 5,972 | 32.1 | 20 | | | 91 | 150 | 247 |
| | Cas9:c-Myc | G1 | 3 | 11,507 | 47.4 | 26 | | | 109 | 162 | 222 |
| Lig4-/- Ku70-/- A1-1 +Ku70 | Cas9:c-Myc | G1 | 1 | 75 | 26.4 | 20 | 23.7 | 3.2 | 39 | 216 | 216 |
| | Cas9:c-Myc | G1 | 2 | 100 | 60.7 | 25 | | | 257 | 313 | 313 |
| | Cas9:c-Myc | G1 | 3 | 156 | 51.2 | 26 | | | 104 | 330 | 841 |
| **Figure 3B** | | | | | | | | | | | |
| Lig4-/- D | Eb-ZFN | G1 | 1 | 279 | 42.3 | 11 | 9.0 | 1.7 | 36 | 679 | 828 |
| | Eb-ZFN | G1 | 2 | 563 | 33.4 | 8 | | | 32 | 568 | 833 |
| | Eb-ZFN | G1 | 3 | 602 | 39.8 | 8 | | | 70 | 638 | 890 |
| Lig4-/- D +Lig4 | Eb-ZFN | G1 | 1 | 9,496 | 5.3 | 4 | 4.0 | 0.0 | 8 | 36 | 545 |
| | Eb-ZFN | G1 | 2 | 8,897 | 4.9 | 4 | | | 8 | 31 | 181 |
| | Eb-ZFN | G1 | 3 | 8,654 | 4.9 | 4 | | | 8 | 29 | 430 |
| Lig4-/- Ku70-/- D1 | Eb-ZFN | G1 | 1 | 15,924 | 18.1 | 12 | 11.3 | 0.6 | 33 | 71 | 256 |
| | Eb-ZFN | G1 | 2 | 15,368 | 17.0 | 11 | | | 31 | 69 | 153 |
| | Eb-ZFN | G1 | 3 | 15,695 | 17.6 | 11 | | | 32 | 71 | 257 |
| Lig4-/- Ku70-/- D1 +Ku70 | Eb-ZFN | G1 | 1 | 905 | 29.6 | 12 | 12.0 | 0.0 | 42 | 384 | 669 |
| | Eb-ZFN | G1 | 2 | 685 | 26.5 | 12 | | | 39 | 423 | 661 |
| | Eb-ZFN | G1 | 3 | 565 | 29.3 | 12 | | | 38 | 460 | 745 |

| | | | | | | | | | | | |
|---|---|---|---|---|---|---|---|---|---|---|---|
| **Figure S4** | | | | | | | | | | | |
| | Cas9:c-Myc | G1 | 1 | 15,747 | 0.6 | 0 | | | 0 | 14 | 54 |
| WT A | Cas9:c-Myc | G1 | 2 | 27,274 | 0.4 | 0 | 0.0 | 0.0 | 0 | 10 | 40 |
| | Cas9:c-Myc | G1 | 3 | 28,610 | 0.4 | 0 | | | 0 | 9 | 27 |
| | Cas9:c-Myc | cycling | 1 | 13,729 | 73.0 | 0 | | | 114 | 1,260 | 4,090 |
| WT A | Cas9:c-Myc | cycling | 2 | 10,678 | 98.4 | 2 | 0.7 | 1.2 | 194 | 1,552 | 8,336 |
| | Cas9:c-Myc | cycling | 3 | 16,013 | 36.0 | 0 | | | 24 | 902 | 3,977 |
| | Cas9:c-Myc | G1 | 1 | 42 | 37.9 | 22.5 | | | 95 | 111 | 111 |
| Lig4-/- A1 | Cas9:c-Myc | G1 | 2 | 51 | 11.5 | 10 | 23.5 | 14.0 | 35 | 43 | 43 |
| | Cas9:c-Myc | G1 | 3 | 36 | 60.1 | 38 | | | 241 | 333 | 333 |
| | Cas9:c-Myc | cycling | 1 | 9,516 | 325.3 | 27 | | | 976 | 3,282 | 9,586 |
| Lig4-/- A1 | Cas9:c-Myc | cycling | 2 | 12,489 | 290.4 | 24 | 24.7 | 2.1 | 909 | 2,763 | 7,833 |
| | Cas9:c-Myc | cycling | 3 | 10,269 | 275.5 | 23 | | | 907 | 3,318 | 8,041 |
| **Figure S6** | | | | | | | | | | | |
| | Cas9:c-Myc | G1 | 1 | 57 | 50.0 | 2 | | | 210 | 512 | 512 |
| Lig4-/- C | Cas9:c-Myc | G1 | 2 | 311 | 67.1 | 24 | 16.7 | 12.7 | 190 | 683 | 758 |
| | Cas9:c-Myc | G1 | 3 | 99 | 81.7 | 24 | | | 292 | 560 | 560 |
| | Cas9:c-Myc | G1 | 1 | 46,294 | 0.7 | 0 | | | 1 | 17 | 49 |
| Lig4-/- C +Lig4 | Cas9:c-Myc | G1 | 2 | 40,169 | 1.1 | 0 | 0.0 | 0.0 | 2 | 21 | 66 |
| | Cas9:c-Myc | G1 | 3 | 43,027 | 0.8 | 0 | | | 2 | 18 | 59 |
| | Cas9:c-Myc | G1 | 1 | 39,069 | 45.3 | 25 | | | 107 | 166 | 341 |
| Lig4-/- Ku70-/- C1 | Cas9:c-Myc | G1 | 2 | 41,770 | 46.9 | 26 | 25.3 | 0.6 | 107 | 163 | 274 |
| | Cas9:c-Myc | G1 | 3 | 38,789 | 45.8 | 25 | | | 107 | 165 | 291 |
| | Cas9:c-Myc | G1 | 1 | 1,936 | 46.9 | 25 | | | 109 | 209 | 763 |
| Lig4-/- Ku70-/- C1 +Ku70 | Cas9:c-Myc | G1 | 2 | 2,922 | 43.4 | 25 | 25.3 | 0.6 | 104 | 157 | 190 |
| | Cas9:c-Myc | G1 | 3 | 1,156 | 48.6 | 26 | | | 112 | 157 | 230 |
| **Figure S8** | | | | | | | | | | | |
| | Eb-ZFN | G1 | 1 | 14,311 | 12.9 | 9 | | | 27 | 56 | 157 |
| Lig4-/- Ku70-/- D2 | Eb-ZFN | G1 | 2 | 14,725 | 17.1 | 11 | 9.7 | 1.2 | 30 | 70 | 256 |
| | Eb-ZFN | G1 | 3 | 5,523 | 12.9 | 9 | | | 27 | 56 | 145 |
| | Eb-ZFN | G1 | 1 | 735 | 26.3 | 12 | | | 31 | 454 | 771 |
| Lig4-/- Ku70-/- D2 +Ku70 | Eb-ZFN | G1 | 2 | 63 | 16.1 | 11 | 11.0 | 1.0 | 33 | 87 | 87 |
| | Eb-ZFN | G1 | 3 | 25 | 13.3 | 10 | | | 31 | 83 | 83 |

* Rejoining junctions are defined as break-site junctions in positive orientation, break-site junctions being defined in Table S1

** Resection range is defined as the maximum length of resection obtained for 90%, 99% or 99.9% of the rejoining junctions, sorted by increasing resection

### Table S3. Jκ1 coding end bait HTGTS-V(D)J-seq libraries information

| | | | Igκv fragments | | | | Igκv12-44 | | | | Igκv16-104 | | | |
|---|---|---|---|---|---|---|---|---|---|---|---|---|---|---|
| **Genotype** | **Rep** | **Reads** | **Total** | **CE** | **RSS** | **CE/RSS Ratio** | **Total** | **CE** | **RSS** | **CE/RSS Ratio** | **Total** | **CE** | **RSS** | **CE/RSS Ratio** |
| **Figure 4A-B (MiSeq)** | | | | | | | | | | | | | | |
| WT A | 1 | 167,151 | 47,130 | 47,031 | 99 | 475.1 | 2,254 | 2,252 | 2 | 1,126.0 | 710 | 709 | 1 | 709.0 |
| | 2 | 167,151 | 52,088 | 51,932 | 156 | 332.9 | 1,792 | 1,792 | 0 | N/A | 450 | 449 | 1 | 449.0 |
| | 3 | 167,151 | 53,074 | 52,880 | 194 | 272.6 | 1,862 | 1,860 | 2 | 930.0 | 566 | 565 | 1 | 565.0 |
| Ku70-/- A1 | 1 | 167,151 | 3,075 | 1,945 | 1,130 | 1.7 | 104 | 71 | 33 | 2.2 | 34 | 20 | 14 | 1.4 |
| | 2 | 167,151 | 5,677 | 4,215 | 1,462 | 2.9 | 235 | 147 | 88 | 1.7 | 54 | 41 | 13 | 3.2 |
| | 3 | 167,151 | 6,383 | 4,477 | 1,906 | 2.3 | 236 | 168 | 68 | 2.5 | 45 | 34 | 11 | 3.1 |
| Lig4-/- A1 | 1 | 167,151 | 35 | 5 | 30 | 0.2 | 0 | 0 | 0 | N/A | 0 | 0 | 0 | N/A |
| | 2 | 167,151 | 37 | 13 | 24 | 0.5 | 3 | 1 | 2 | 0.5 | 1 | 1 | 0 | N/A |
| | 3 | 167,151 | 74 | 57 | 17 | 3.4 | 2 | 1 | 1 | 1.0 | 1 | 1 | 0 | N/A |
| Lig4-/- A1 +Lig4 | 1 | 167,151 | 11,309 | 11,083 | 226 | 49.0 | 530 | 514 | 16 | 32.1 | 183 | 182 | 1 | 182.0 |
| | 2 | 167,151 | 10,577 | 10,278 | 299 | 34.4 | 538 | 519 | 19 | 27.3 | 184 | 182 | 2 | 91.0 |
| | 3 | 167,151 | 13,708 | 13,179 | 529 | 24.9 | 667 | 621 | 46 | 13.5 | 234 | 231 | 3 | 77.0 |
| **Figure 4C-D (MiSeq)** | | | | | | | | | | | | | | |
| Lig4-/- A1 | 1 | 167,151 | 17 | 6 | 11 | 0.5 | 0 | 0 | 0 | N/A | 0 | 0 | 0 | N/A |
| | 2 | 167,151 | 32 | 13 | 19 | 0.7 | 0 | 0 | 0 | N/A | 0 | 0 | 0 | N/A |
| | 3 | 167,151 | 65 | 54 | 11 | 4.9 | 1 | 1 | 0 | N/A | 2 | 2 | 0 | N/A |
| Lig4-/- Ku70-/- A1-1 | 1 | 167,151 | 1,867 | 1,100 | 767 | 1.4 | 48 | 30 | 18 | 1.7 | 23 | 5 | 18 | 0.3 |
| | 2 | 167,151 | 3,419 | 2,289 | 1,130 | 2.0 | 142 | 78 | 64 | 1.2 | 42 | 28 | 14 | 2.0 |
| | 3 | 167,151 | 4,188 | 2,650 | 1,538 | 1.7 | 158 | 101 | 57 | 1.8 | 55 | 42 | 13 | 3.2 |
| Lig4-/- Ku70-/- A1-1 +Ku70 | 1 | 167,151 | 46 | 34 | 12 | 2.8 | 0 | 0 | 0 | N/A | 1 | 1 | 0 | N/A |
| | 2 | 167,151 | 75 | 40 | 35 | 1.1 | 5 | 3 | 2 | 1.5 | 2 | 2 | 0 | N/A |
| | 3 | 167,151 | 60 | 37 | 23 | 1.6 | 2 | 2 | 0 | N/A | 0 | 0 | 0 | N/A |
| **Figure S9A-B (NextSeq)** | | | | | | | | | | | | | | |
| WT B | 1 | 600,000 | 291,174 | 290,303 | 871 | 333.3 | 13,215 | 13,168 | 47 | 280.2 | 5,433 | 5,428 | 5 | 1,085.6 |
| | 2 | 600,000 | 292,983 | 292,146 | 837 | 349.0 | 13,437 | 13,416 | 21 | 638.9 | 5,316 | 5,300 | 16 | 331.3 |
| | 3 | 600,000 | 305,846 | 304,972 | 874 | 348.9 | 13,816 | 13,778 | 38 | 362.6 | 5,480 | 5,470 | 10 | 547.0 |
| Ku70-/- B1 | 1 | 600,000 | 50,218 | 35,292 | 14,926 | 2.4 | 2,059 | 1,326 | 733 | 1.8 | 1,106 | 819 | 287 | 2.9 |
| | 2 | 600,000 | 59,967 | 41,837 | 18,130 | 2.3 | 2,421 | 1,565 | 856 | 1.8 | 1,337 | 983 | 354 | 2.8 |
| | 3 | 600,000 | 60,811 | 42,063 | 18,748 | 2.2 | 2,406 | 1,536 | 870 | 1.8 | 1,375 | 1,015 | 360 | 2.8 |
| Lig4-/- B1 | 1 | 600,000 | 50 | 34 | 16 | 2.1 | 4 | 2 | 2 | 1.0 | 1 | 0 | 1 | N/A |
| | 2 | 600,000 | 55 | 38 | 17 | 2.2 | 6 | 3 | 3 | 1.0 | 2 | 2 | 0 | N/A |
| | 3 | 600,000 | 60 | 42 | 18 | 2.3 | 2 | 2 | 0 | N/A | 1 | 0 | 1 | N/A |
| **Figure S9C-D (MiSeq)** | | | | | | | | | | | | | | |
| Lig4-/- C | 1 | 377,068 | 16 | 12 | 4 | 3.0 | 1 | 1 | 0 | N/A | 1 | 0 | 1 | N/A |
| | 2 | 377,068 | 3 | 3 | 0 | N/A | 0 | 0 | 0 | N/A | 0 | 0 | 0 | N/A |
| | 3 | 377,068 | 46 | 46 | 0 | N/A | 0 | 0 | 0 | N/A | 0 | 0 | 0 | N/A |
| Lig4-/- Ku70-/- C1 | 1 | 377,068 | 4,457 | 3,045 | 1,412 | 2.2 | 186 | 117 | 69 | 1.7 | 33 | 20 | 13 | 1.5 |
| | 2 | 377,068 | 5,874 | 3,902 | 1,972 | 2.0 | 249 | 159 | 90 | 1.8 | 50 | 42 | 8 | 5.3 |
| | 3 | 377,068 | 7,225 | 5,172 | 2,053 | 2.5 | 360 | 253 | 107 | 2.4 | 71 | 47 | 24 | 2.0 |
| Lig4-/- Ku70-/- C1 +Ku70 | 1 | 377,068 | 436 | 304 | 132 | 2.3 | 17 | 8 | 9 | 0.9 | 1 | 1 | 0 | N/A |
| | 2 | 377,068 | 612 | 372 | 240 | 1.6 | 14 | 9 | 5 | 1.8 | 0 | 0 | 0 | N/A |
| | 3 | 377,068 | 589 | 370 | 219 | 1.7 | 29 | 20 | 9 | 2.2 | 3 | 0 | 3 | N/A |
| Lig4-/- C +Lig4 | 1 | 377,068 | 35,059 | 34,932 | 127 | 275.1 | 1,048 | 1,042 | 6 | 173.7 | 514 | 513 | 1 | 513.0 |
| | 2 | 377,068 | 20,882 | 20,813 | 69 | 301.6 | 882 | 879 | 3 | 293.0 | 233 | 232 | 1 | 232.0 |
| | 3 | 377,068 | 25,600 | 25,521 | 79 | 323.1 | 1,034 | 1,028 | 6 | 171.3 | 324 | 324 | 0 | N/A |

**Table S4. Primers and oligos used in this study**

| | |
|---|---|
| HTGTS Primers | |
| Jk1 bio primer | /5Bio/TTCCCAGCTTTGCTTACGGAG |
| Jk1 nested primer (MiSeq) | AGTGCCAGA ATCTGGTTTCAGAG |
| Jk1 nested primer (NextSeq) | CAGACATAGACAACGGAAGAAAG |
| *c-Myc* bio primer | /5Bio/GCCTCGGCTCTTAGCAGACTG |
| *c-Myc* nested primer | CCTCGGCTCTTAGCAGACTG |
| EB bio primer | /5Bio/CAGAAGCCTTCAGTATGCACCA |
| EB nested primer | TGAGTCACAGGAACAGAGTC |
| sgRNAs | |
| Cas9-*c-Myc* F | CACCGGACGAGCGTCACTGATAGTA |
| Cas9-*c-Myc* R | AAACTACTATCAGTGACGCTCGTCC |
| Cas9-sgRNA1 F (Ku70 knockout) | CACCGGCAGTGCAACACTGGTTCCA |
| Cas9-sgRNA1 R (Ku70 knockout) | AAACTGGAACCAGTGTTGCACTGCC |
| Cas9-sgRNA2 F (Ku70 knockout) | CACCGGATGGTGCCAGGCCAGATGT |
| Cas9-sgRNA2 R (Ku70 knockout) | AAACACATCTGGCCTGGCACCATCC |
| Genotyping | |
| Ku70_F (Ku70 knockout screening) | CCTCCAGGGCTATGTTCGAA |
| Ku70_R (Ku70 knockout screening) | CCTGGCTTCTTCAGATGCAT |
| Ku70_FgI (Ku70 knockout screening) | GCAGGCCTCAAGCAGGCCAT |
| Southern probe | |
| Ku70 probe (knockout screening) | CTGGTAGGTGGCTAACCTTTCCTACCGAATCTTGTTTAAGAACTG ACAGAGAGGTGGGGCTGTAGAGGTGGCTCAGCAGTTAAGAGAA CATATCCTGTCCATGCATCGTACCCAGGTTCTGGTCCCAGCATC CACATGGTGGCTCACAGCCAACCATACTTTGGTTTCAGGAGACT AGACCCCTTCTGACCTCTGCATGCATGAGGCATGCACGAGGCG TGCACATGCTGCACATACATACATGCAGATAAATAAATCTTTAAAC AACAGGAGCTGGACATGATGGCACATGACTTTAATCTCAGAACT CAGGAGACAAAAGCAGGCAGATCTCTGGGGACAGTCTGGGTTA CAAGAGTTCGTTCCAGGACAGACAGGACTGTGTAGAGACCATG TCTGGGCGAAAGAAAACCAGACAGACACACTGAGCAAGCTAGC CAGTGAAGTAAGTCACTGTAAACTGCGTCTTTCTGGATAAACGG TCAGGACAGGAAGCAGCTGGCCTTGCA |
| *c-Myc* probe | GACGACGAGACCTTCATCAAGAACATCATCATCCAGGACTGTAT GTGGAGCGGTTTCTCAGCCGCTGCCAAGCTGGTCTCGGAGAA GCTGGCCTCCTACCAGGCTGCGCGCAAAGACAGCACCAGCCT GAGCCCCGCCCGCGGGCACAGCGTCTGCTCCACCTCCAGCCT GTACCTGCAGGACCTCACCGCCGCCGCGTCCGAGTGCATTGAC CCCTCAGTGGTCTTTCCCTACCCGCTCAACGACAGCAGCTCGC |

| | |
|---|---|
| | CCAAATCCTGTACCTCGTCCGATTCCACGGCCTTCTCTCCTTCC TCGGACTCGCTGCTGTCCTCCGAGTCCTCCCCACGGGCCAGC CCTGAGCCCCTAGTGCTGCATGAGGAGACACCGCCCACCACCA GCAGCGACTCTGGTAAGCTACCCCATTCACAGCAGGGTAGGAA GCGAGAGGTTGGATGGACCTCCTTCTCCACCACTCATTGGCATT AATTCAATTGGCCTCCGGGGCTCCCCCTTTCTTTCCCTTCTGTC TAAGAGCTCTTCATCCCTGGATTC |
| Jk3 probe (Bredemeyer et al., 2006) | TTCGGTGCTGGGACCAAGCTGGAGCTGAAACGTAAGTACACTT TTCTCATCTTTTTTTATGTGTAAGACACAGGTTTTCATGTTAGGAG TTAAAGTCAGTTCAGAAAATCTTGAGAAAATGGAGAGGGCTCAT TATCAGTTGACGTGGCATACAGTGTCAGATTTTCTGTTTATCAAG CTAGTGAGATTAGGGGCAAAAAGAGGCTTTAGTTGAGAGGAAAG TAATTAATACTATGGTCACCATCCAAGAGATTGGATCGGAGAATA AGCATGAGTAGTTATTGAGATCTGGGTCTGACTGCAGGTAGCGT GGTCTTCTAGACGTTTAAGTGGGAGATTTGGAGGGGATGAGGA ATGAAGGAACTTCAGGATAGAAAAGGGCTGAAGTCAAGTTCAGC TCCTAAAATGGATGTGGGAGCAAACTTTGAAGATAAACTGAATGA CCCAGAGGATGAAACAGCGCAGATCAAAGAGGGGCCTAGAGCT |
| ERag probe (Tubbs et al., 2014) | AACTTCCTCCAGCAGGCGATCTGCCAGTTGCAAGAGTATCAAAA CAATGCTAAGCCCTCGGGTTCTGTCTAATACTCGACAGAAAATAA CTGCGCAGGGCTACACAGTGAGGATTATTTTTCTTGCTGTGTTTT CTTTTATGAAAGAAAGCTGTTTTTCTGCTCGACTGAAATTCCTCT CCTGCTTGGAAATGGGTAAGAGGGCCAGCTGCTTGCTATATTTT TCTCTGTGGTATTTTTCTACAGCAGATTTTAAAATTCATTCTTCAG ACTGGGAAAAGCCTCTCTTTGTACCCAACCTCACTCGTGAAAGT TTAAAGATTCATTTCTAACGCTCAAATTCAAAATAGCTTTGGGGA CCAAAATATGTCAGCTTTAGAAACATGTTGTTTTATGTGCTGTCTC CAGTTTCATTTTTAATTACATTTTTTAGAAGTTGTCAAGTATGAGTA GTAATATTCCTTTGGCTGAACTGACAGTCAA |
| Eb probe (Lee et al., 2013) | GTTAACCAGGCACAGTAGGACCCTTAATACTTGTTTCCCTGACC GATTCCAGCAAAGAGATATAGGAACATGATAAAAAAAATAATCTTA AGAAGAACTGTTTTTCATCAAACATCAGCAAGTGAAGAGATGCAT TCCTGGGACTTTTCGGTTCCTGAAGACAATGGGGGAAGGGGTG GAAGCATCTCACCCCAGGTCTGGCTGTTTATCTGTAAGTAACATC AGCACCAAGTAAGAATGGCCACCTGCCATAGCTCCATCTCCAGG AGTCACAACAGGATGTGGTTTGACATTTACCAGGTCCTACATCT GGGGTGCCTGTGAATGCTCCCCCACTCACTCACATTCTGAGCAT TTTGGGAACCACACTTGCCACATCCTGTCTTCAAACCCTTCTCA TGCAGCCCTTTCTACCTCAGCCTCTAGCTATCACCGTGACAGTG AGAACAACCAAGAGGAGGGTCTCTAAAGGGGAACCCATATATCT CATCTCAAGCCCCCCATCTAAACTATCTTACATCTGTTAGAAGTC ACACAGGAAACAGAAGCCTTCAGTATGCACCATGG |
| 53bp1 probe | TAGAATGTGAGCCCCCTCTGCCCATGGATGGAGTTGAAAGCTGT AAACTCAGGGCTCTCCCACGAAGCTACATCCTCAGCTCTGATTA TGTTTTTTGTCTTCATTTTTCCAAACCAATTTTAGTTATTTTTGCTT ATAATTTTCTGTTTCCATGGATACTTAGAAAAGACCTAATGTTAAT GATGTGCCCAGAATGGCTCCTGCCTGCTCCTCTCCGTAACCTCC TTAGCTCTCTTCTCTCCACAGGGAACTCCTTCTGTCTTCTAGGTT AAAGTCCTTGCTAGTAGCTTATGCACTTTCCCAGTAGAAAGA |
